# Localized mRNAs and protein synthesis in cortical layer 1

**DOI:** 10.1101/2025.11.23.689989

**Authors:** Teresa Spanò, Belquis Nassim-Assir, Eva Kaulich, Georgi Tushev, Nicole Fürst, Helene Will, Lucija Marić, Elena Ciirdaeva, Erin M. Schuman

## Abstract

Cortical layer 1 plays an essential role for brain function, integrating many streams of information and undergoing synaptic plasticity during learning. Yet, we understand little of the underlying molecular processes, and whether they are specialized to achieve layer-specific functions. Here, we show that layer 1 and its synapses are metabolically active. Using laser capture RNA-sequencing and fluorescence *in situ* hybridization, we report an abundance of synaptic transcripts localized to layer 1. We purified layer 1 excitatory and inhibitory synapses and characterized their enriched transcriptomes, identifying candidate proteins that can be locally synthesized to maintain and regulate synaptic transmission in layer 1. Furthermore, we find significant differences between the transcriptome of synapses in layer 1 and in deeper, somatic layers, suggesting that local translation might confer specialized function. Finally, by comparing our tissue data with the transcriptome from the hippocampal CA1 strata, we discover a strong similarity between cortical layer 1 and stratum lacunosum moleculare, suggesting that these distal layers share a common molecular environment. Together, our results establish that local protein synthesis is an important mechanism for layer 1, and provide the first comprehensive characterization of the transcripts localized to layer 1 and its synapses.

## Introduction

The mammalian neocortex is responsible for processing sensory information, planning and executing movements, and integrating multiple senses to create a unified understanding of the world. It contains 6 layers with different cell-types, projections and synapses, resulting in layer-specific synaptic properties, computations and functions ^1,2^. Layer 1, at the pia surface, differs from the other layers because of the paucity of neuronal cell bodies ^3,4^. Rather, the layer 1 space is predominantly occupied by neuropil, comprising the apical dendrites of pyramidal cells, a wide diversity of axonal projections arising from higher cortical areas, memory centers and neuromodulatory nuclei, and a high density of synapses ^5–7^. Integrating this large variety of inputs, layer 1 is thought to play a crucial role for many higher cortical functions, including learning, attention and predictive coding ^4,8,9^. Importantly, layer 1 is a highly plastic area, displaying learning-related changes in excitatory and inhibitory inputs ^10–14^, apical dendrites and spines ^15,16^, and in local, layer 1 inhibitory interneurons ^17^. Layer 1 synapses also display specific forms of plasticity ^18–22^. While recent work has focused on understanding layer 1 computations and plasticity, the underlying molecular processes – and how they differ from other cortical layers – remain largely unexplored.

Decades of work have shown that neurons localize mRNAs to their dendrites and axons and use local protein synthesis to maintain and regulate their synaptic proteomes with high spatial and temporal resolution ^23^. This process is required for many forms of long-term synaptic plasticity ^24–26^, as well as for the maintenance of synaptic transmission at certain synapses ^27,28^. Recent data indicate that hippocampal synapses are characterized by different transcriptomes depending on their location within the dendritic tree, suggesting that local protein synthesis contributes to the molecular specialization of different synapses within the same neurons ^29^. While most work on local protein synthesis in the mammalian brain has been carried out in hippocampal slices or dissociated cultures, there is evidence that mRNAs are localized and translated also at cortical synapses ^30–33^. However, it is unknown how local protein synthesis is regulated across cortical layers and if the synapses that are most remote from cell bodies, like those in layer 1, exhibit a unique profile of mRNA localization and translation.

Here, we establish that protein synthesis is a feature of rodent neocortical layer 1, its neuropil and its synapses. Using laser capture RNA-sequencing, we characterize the layer 1 transcriptome, which includes many synaptic transcripts. By purifying excitatory and inhibitory synapses from layer 1, we then describe their enriched transcriptomes, comprising more than 1000 and 150 transcripts, respectively, and identify significant differences to the transcriptomes of synapses in deeper cortical layers. As one example, we found that the transcript encoding the synaptic vesicle protein Complexin 3 is locally translated in layer 1, and that there is a corresponding layer 1 enrichment of the protein. Finally, we compare our cortical data to the strata in the hippocampus, discovering a transcriptomic similarity between neocortical layer 1 and stratum lacunosum moleculare, which is also found in the molecular layer of other 3-layered cortices.

## Results

### Layer 1 neuropil and synapses are rich in localized mRNAs and actively synthesize proteins

To investigate the potential for local protein synthesis in cortical layer 1, we first examined whether there is a population of localized mRNAs. Using fluorescent *in situ* hybridization (FISH) against the *polyA* sequence common to all mRNAs, we visualized mRNA distribution in layer 1 of coronal cortical slices from adult mice. As expected, *polyA* FISH signal labelled layer 1 cell bodies (mostly from non-neuronal cells; Figure S1A), but there were also puncta present throughout the neuropil. In contrast, almost no puncta were observed when using the complementary control probe, *polyT* (Figure 1A). To quantify the mRNA density in neuropil-rich areas, we digitally excluded the regions marked by the nuclear signal of 4′,6-diamidino-2-phenylindole (DAPI), then quantified the FISH signal revealing an 8-fold increase in puncta density with the *polyA* probe (Figure 1B).

**Figure 1.**
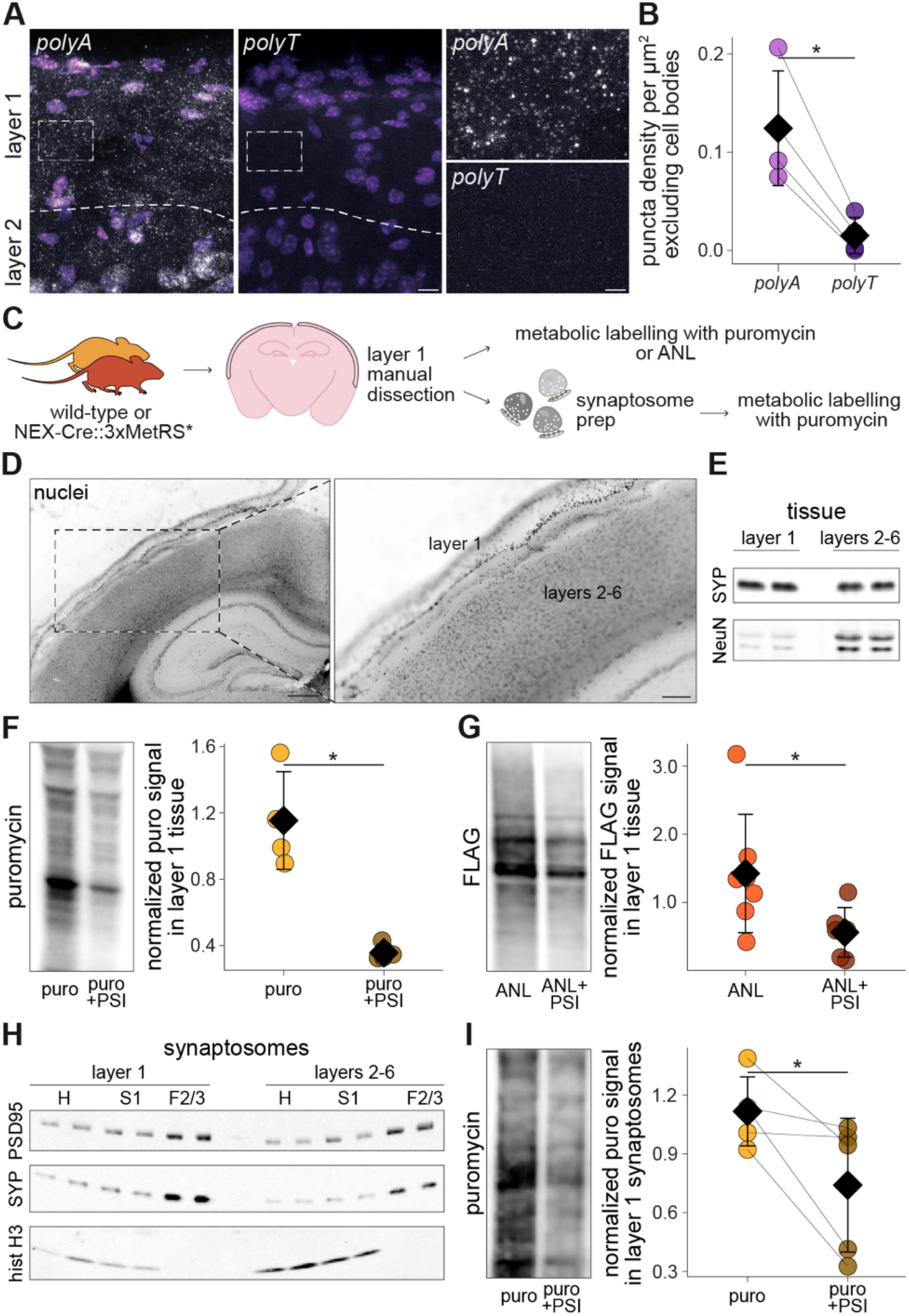
Layer 1 neuropil and synapses are rich in localized mRNAs and actively synthesize proteins. A) Representative mouse cortex FISH images using *polyA* or *polyT* probes (grey); nuclei are stained with DAPI (purple). Dashed lines indicate the layer 1-2 boundary. Dashed rectangles indicate an area devoid of cell bodies, shown at higher magnification on the right. Scale bar on the left and right = 10 μm and 3 μm, respectively. B) Density of FISH puncta in layer 1 after digital exclusion of cell bodies (n = 4 brains, paired t-test). C) Pipeline for metabolic labelling experiments: acute slices from wild-type or NEX-Cre::3xMetRS* mice were manually dissected to isolate layer 1 (panels D and E); the tissue was labelled with either puromycin (F) or ANL (G), or it was used for synaptosome preparations (panel H), which were labelled with puromycin (I). D) Representative image of a layer 1 dissection from a coronal slice: overall view (left) and zoom in on layer 1 (right). Scale bar on the left and right = 500 and 200 μm, respectively. E) Representative Western blot from layer 1 and respective layers 2-6 from two brains, stained for synaptophysin (SYP; synapses) and NeuN (neuronal cell bodies). F) Representative Western blot (left) and quantification (right) of protein synthesis in dissected layer 1 tissue (n = 5 brains, Welch Two Sample t-test). Slices were treated with puromycin (puro; 18 μM, 15 min) +/- the protein synthesis inhibitor emetine (180 μM). G) Representative Western blot (left) and quantification (right) of protein synthesis in dissected layer 1 tissue from NEX-Cre::3xMetRS* mice (n = 6 or 7 brains, Welch Two Sample t-test). Slices were labelled with ANL (1 mM, 1 hour) +/- the protein synthesis inhibitor cycloheximide (355 μM). H) Representative Western blot of different fractions of the synaptosome preparation from layer 1 and layers 2-6 dissected tissues from two mice, stained for the synaptic proteins PSD95 and synaptophysin (SYP), and the nuclear histone H3. I) Representative Western blot (left) and quantification (right) of protein synthesis in synaptosomes derived from dissected layer 1 tissue (synaptosomes split into two conditions, n = 5 brains, paired t-test). Synaptosomes were treated with puro (240 μM, 15 min) +/- the protein synthesis inhibitor emetine (4 mM). * p < 0.05. Points represent individual observations; the black diamond and the error bars represent the mean and standard deviation. For statistical details, see Supplementary Table 1.

The abundance of mature mRNAs throughout layer 1 suggests the possibility of local translation. To address this, we performed metabolic labelling experiments, initially using puromycin ^34^, on manually microdissected layer 1 tissue, processing layers 2-6 in parallel (Figure 1C-D; see methods). As expected, relative to layers 2-6, layer 1 was depleted of cell bodies (labelled by NeuN) but contained more synaptophysin, reflecting a higher synaptic density (Figure 1E, S1B). To label nascent protein synthesis, microdissected tissue was treated with puromycin (15 min), lysates were prepared and nascent proteins were detected with an anti-puromycin antibody. In layer 1 there was a strong nascent protein signal that was sensitive to treatment with the protein synthesis inhibitor (PSI) emetine (Figure 1F, S1C); a similar signal was observed in layers 2-6 (Figure S1D). The nascent protein signal detected in layer 1 could, in principle, arise from neuronal axons and dendrites or non-neuronal cells that reside in the area. To visualize directly the nascent proteins present in neocortical excitatory neurons, we made use of a mouse line where bioorthogonal labeling of nascent proteins can be achieved in specific cell-types via the expression of a mutant methionyl-tRNA synthetase (MetRS*) and the application of a non-canonical amino acid (NCAA) followed by click chemistry ^35^ (Figure 1C). Cortical layer 1 (or layers 2-6) was microdissected from NEX-cre::3xMetRS* mice, treated with the NCAA azidonorleucine (ANL) for 1 hour, and then click chemistry with a FLAG tag was performed. Again, we detected a nascent protein signal in layer 1 that was significantly inhibited by a PSI (Figure 1G, S1E); the same pattern was observed in layers 2-6 (Figure S1F).

To test whether some of the above protein synthesis signal may arise from synapses, we prepared synaptosomes from microdissected layer 1 tissue and, as a control, from layers 2-6. Despite the low amount of starting material, we found that layer 1 synaptic proteins were enriched relative to the input homogenate at a level equivalent to that observed in the layers 2-6 synaptosomes (Figure 1H, S1G). Isolated layer 1 (and layers 2-6) synaptosomes were then briefly (15 min) metabolically labelled with puromycin and nascent proteins were detected using anti-puromycin antibodies, revealing a robust signal that was significantly inhibited by a PSI (Figure 1I, S1H-I). These results indicate that there is ongoing protein synthesis at synapses throughout the cortical layers, including in layer 1.

### Synaptic transcripts are abundant in cortical layer 1

To identify the specific transcripts present in layer 1 we used laser capture microdissection to collect cortical layer 1 (Figure 2A) and, in parallel, the corresponding somatic layers (layers 2-6). We extracted RNA from all samples and processed it for RNA-sequencing (RNA-seq) yielding greater than 17 million reads per replicate, with greater than 70% uniquely mapped to the mouse genome (Figure S2A). We detected 17447 and 17114 genes in layer 1 and layers 2-6, respectively; a principal component analysis (PCA) of the 500 most variable genes revealed that 89% of the variation mapped onto the component that separated the layer 1 samples from layers 2-6 (Figure 2B). Differential expression (DE) analysis revealed 3200 and 2838 transcripts enriched in layer 1 and layers 2-6, respectively (Figure 2C). The top gene ontology (GO) terms associated with the genes enriched in layer 1 included “extracellular matrix”, “angiogenesis” and “cell-cell junction”, reflecting the vicinity of layer 1 to pia structures (Figure 2D). Conversely, the transcripts associated with the genes enriched in the cell body layers were related to synaptic assembly, synaptic vesicles, and ion transporters (Figure S2B). This is to be expected, as in layer 1 most cell bodies, a major source of mRNAs, belong to non-neuronal cells, while deeper layers are densely packed with neuronal somata. Indeed, we found that markers for astrocytes, glia limitans and endothelial cells were enriched in layer 1 (Figure S2C). *Ndnf* and *Reelin*, two markers for layer 1 interneurons, were also enriched in layer 1, while markers for neuronal somata, axon initial segments, deeper layer interneurons and layer 4 neurons were enriched in layers 2-6 (Figure S2C). We then looked at synaptic transcripts in layer 1 using SynGO (www.syngo.org)^36^, and the transcripts found at cortical synapses in SynDive (www.syndive.org)^37^. Compared to the total layer 1 transcriptome, these transcripts were expressed at higher level (Figure 2E, S2D), suggesting that mRNAs encoding synaptic proteins are abundant in layer 1. Indeed, 71 of the 200 most abundant transcripts in this layer were listed in SynGO, and 30 encoded proteins found in cortical synapses (Figure 2F). Among the most highly expressed layer 1 synaptic mRNAs were the three isoforms of Calmodulin (*Calm1/2/3*), their associated kinase *Camk2a*, their binder *Nrgn* and several ribosomal protein genes. To validate the presence of a subset of synaptic transcripts in layer 1, we performed FISH with probes against the *Camk2a*, *Ddn, Kif5a*, *Shank1, Actb* and *Dlg4* mRNAs along with no probe controls. For all candidate transcripts, FISH puncta were present throughout the layer 1 neuropil; the puncta density was significantly higher than the negative control (Figure 2G, S2E-F). Overall, this first comprehensive transcriptome of cortical layer 1 tissue revealed that transcripts encoding synaptic proteins are abundant in this distal layer.

**Figure 2.**
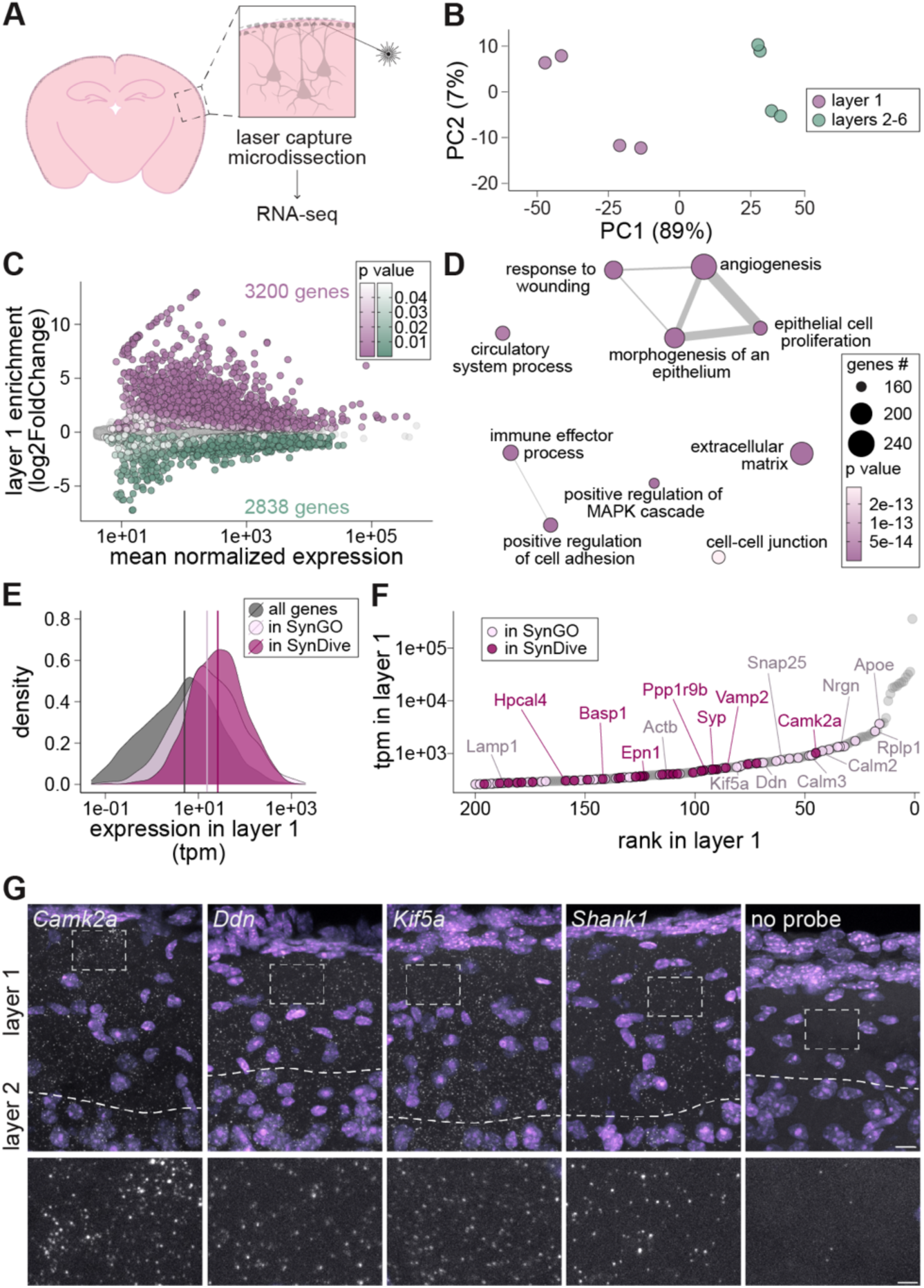
Synaptic transcripts are abundant in cortical layer 1. A) Schematic of layer 1 laser capture microdissection. B) First two components in the principal component analysis of the 500 most variable genes. C) Mean normalized expression of genes against their log2FoldChange for layer 1 versus deeper layers: highlighted are the 3200 and 2838 genes that were enriched in layer 1 and layers 2-6, respectively (no log2FoldChange cut-off, adjusted p value < 0.05). D) The 10 most overrepresented GO terms associated with the genes enriched in layer 1. The terms are ranked by gene count. The thickness of the edge represents similarity between the terms (only above 0.15). E) Density plots showing the distribution of expression of all genes in layer 1 (grey) and of the subsets corresponding to genes listed in SynGo (light pink) and in SynDive (purple, representing the union of proteins found at cortical inhibitory and excitatory synapses). F) The 200 transcripts with the highest expression in layer 1 based on transcripts per million (tpm). Genes listed in SynGO and in SynDive are highlighted in light pink and purple, respectively. G) Representative mouse cortex FISH images using probes against the indicated synaptic transcripts (grey), with nuclei stained with DAPI (purple). Dashed lines indicate the layer 1-2 boundary. Dashed rectangles indicate an area devoid of cell bodies, shown at higher magnification on the bottom. Scale bar on the top and bottom = 10 and 3 μm, respectively.

### The transcriptome of excitatory synapses in layer 1

The analysis above shows that synaptic transcripts are abundant in layer 1, but does not indicate their cellular or subcellular origin. To directly determine the transcriptomes of layer 1 synapses, we prepared synaptosomes from dissected layer 1 (and layers 2-6 as control) from a mouse line in which excitatory synapses were labelled with tdTomato (see methods). Using fluorescence-activated synaptosome sorting ^33,37,38^, we enriched for layer 1 and layers 2-6 excitatory synapses, reaching > 90% purity (tdTom+ fraction; Figure 3A, S3A). As a negative control, we sorted particles (from the same starting material) of a similar size that were tdTomato negative (non-excitatory synapses and contaminants, tdTom- fraction). All samples were processed for RNA-seq, yielding more than 2 million reads (Figure S3B). A PCA revealed clear clustering according to layer and fraction, with the unsorted synaptosomes closely resembling the tdTom+ fraction, as expected (most of the cortical synaptosomes are excitatory; Figure 3B, S3A). DE analysis of the layer 1 tdTom+ synapses and the controls revealed 1052 genes enriched in layer 1 excitatory synapses (Figure 3C), including *Camk2a, Ddn*, *Kifc2* and *Ncs1.* The top 10 GO terms associated with the enriched transcripts were related to synaptic organization and modulation, protein kinase and ion transport, suggesting that the transcriptome of layer 1 excitatory synapses encodes molecules that modulate synaptic transmission and signaling (Figure 3D).

**Figure 3.**
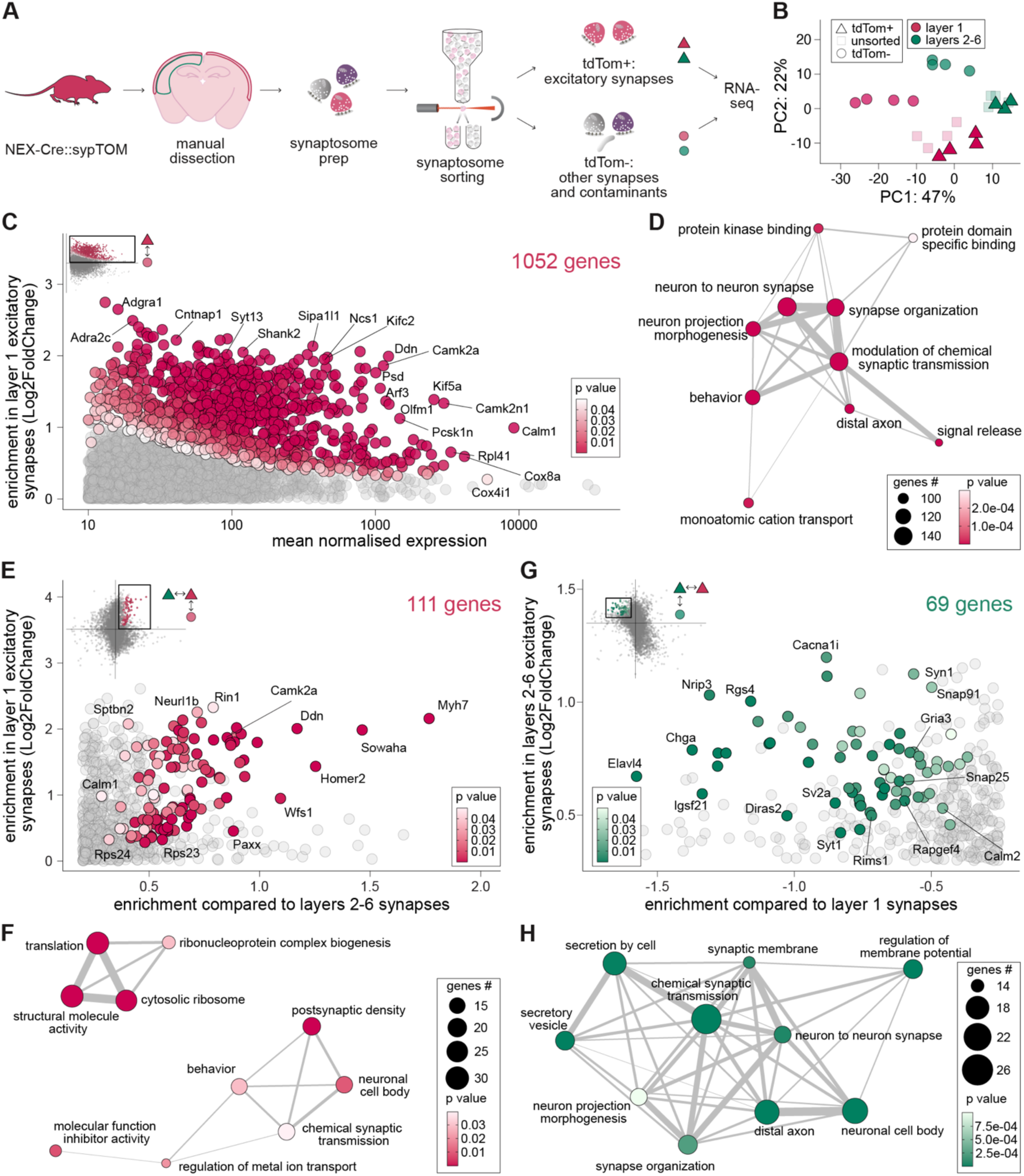
The transcriptome of layer 1 excitatory synapses and its comparison to layers 2-6. A) Pipeline for sorting excitatory synapses from layer 1. B) First two components in the principal component analysis of the 500 most variable genes. C) Mean normalized expression of genes comparing layer 1 tdTom+ fraction (excitatory synapses) to the tdTom- (negative control): highlighted are the 1052 genes that were enriched in the synapses (no log2FoldChange cut-off, adjusted p value < 0.05). D) The 10 most overrepresented GO terms associated with the genes enriched in C. The terms are ranked by gene count. Thickness of the edge represents similarity between the terms (only above 0.15). E) Scatterplot comparing the tdTom+ fraction of layer 1 and that of layers 2-6 (x axis) plotted against the layer 1 of tdTom+ fraction against its negative control (y axis). Highlighted are the 111 transcripts that were significantly enriched in excitatory synapses of layer 1 in both comparisons. F) The 10 most overrepresented GO terms associated with the genes enriched in E. The terms are ranked by gene count. Thickness of the edge represents similarity between the terms (only above 0.15). G) Scatterplot of log2FoldChange comparing the tdTom+ fraction of layer 1 and that of layers 2-6 (x axis) against the log2FoldChange in layers 2-6 of tdTom+ fraction against its negative control (y axis). Highlighted are the 69 transcripts that are significantly enriched in layers 2-6 synapses in both comparisons. H) The 10 most overrepresented GO terms associated with the genes enriched in G. The terms are ranked by gene count. Thickness of the edge represents similarity between the terms (only above 0.15).

Next, we compared the transcriptome of layer 1 and layers 2-6 tdTom+ fractions. We identified 111 genes that were significantly enriched in layer 1 excitatory synapses, both compared to layers 2-6 excitatory synapses and to their layer 1 tdTom- control (Figure 3E). Within this set, a group of transcripts was related to postsynaptic function and behavior, encoding many synaptic proteins including scaffolding proteins (*Shank2*, *Homer2*, *Cnksr2*), signaling modulators (*Arhgap33*, *Git1, Camk2a, Camk2n1, Ncs1*) and receptor and channel auxiliary proteins (*Cnih2, Cacnb3*; Figure 3F). A second group was related to translation, with many transcripts encoding either ribosomal proteins or ribosome-interacting proteins (Figure 3F). Conversely, we identified 69 genes that were de-enriched in layer 1 synapses but significantly enriched in synapses of other layers (Figure 3G). Among these, many encoded channel subunits or auxiliary proteins leading to GO terms such as “regulation of membrane potential” and “synaptic membrane” (Figure 3H). Additionally, we found many transcripts encoding synaptic vesicle proteins (*Rims1, Snap25*, *Snap91*, *Stxbp1*, *Synj1*, *Syt1*; Figure 3G). Overall, these results indicate that the excitatory synapses of layer 1 and layers 2-6 have different transcriptomic signatures, potentially enabling layer-specific functions.

### Layer 1 inhibitory synapses contain a distinct set of localized mRNAs

We next determined the transcriptome of inhibitory synapses in layer 1 using the above described pipeline with a GAD2-Cre driver line^37^ (Figure 4A). Visual inspection of these mice revealed that tdTomato was also expressed in a small subset of astrocytes (Figure S4A). A suite of control experiments revealed that there was no astrocytic contamination in the downstream synaptosome preparation (Figure S4B-D). We thus sorted tdTom+ particles from microdissected layer 1 and layers 2-6 tissue, reaching ∼87% and 80% in layer 1 and 2-6, respectively (Figure S5A). RNA-seq yielded more than 2 million reads per sample, with two exceptions that were excluded from further analysis (Figure S5B). After correcting for individual brain variation, a PCA showed clear clustering between the layers, and also between sorted synaptosomes and control fractions (Figure 4B, Figure S5C). DE analysis of the tdTom+ and tdTom- fractions of layer 1 resulted in 182 genes enriched in layer 1 inhibitory synapses (Figure 4C). With the highest fold change (> 4), we found the transcript encoding the alpha 4 nicotinic receptor subunit (*Chrna4*). Among the most highly expressed and significantly enriched were many transcripts encoding transport proteins (*Kif5a, Kif5b, Kif5c, Kif1a, Klc1*) and several synaptic modulators, resulting in GO terms related to synaptic transmission, membrane potential and trafficking (Figure 4D). Many enriched transcripts also encoded signaling proteins, such as tyrosine kinases and proteases, and immune system players (Figure 4C-D). When comparing the inhibitory synaptosomes of layer 1 and layers 2-6, 38 genes were enriched in layer 1 (Figure 4E). These included the genes encoding the proteins Reelin and the Src family tyrosine kinase Lyn, the complement proteins C1qc and C1qb, the purinergic receptor subunit P2ry12, the vesicle exocytosis protein Complexin 3 and the transcription factor COUP-TFII (*Nr2f2*). Only 12 transcripts were de-enriched in layer 1 but enriched layers 2-6 inhibitory synapses (Figure 4F). These transcripts encoded, among others, the transport protein Kif5b and the kinesin light chain 1 (*Klc1*), the calcium-binding protein Parvalbumin, the vesicle exocytosis protein Complexin 1 and two guanine nucleotide exchange factors (*Rapgef4*, *Mcf2l*).

**Figure 4.**
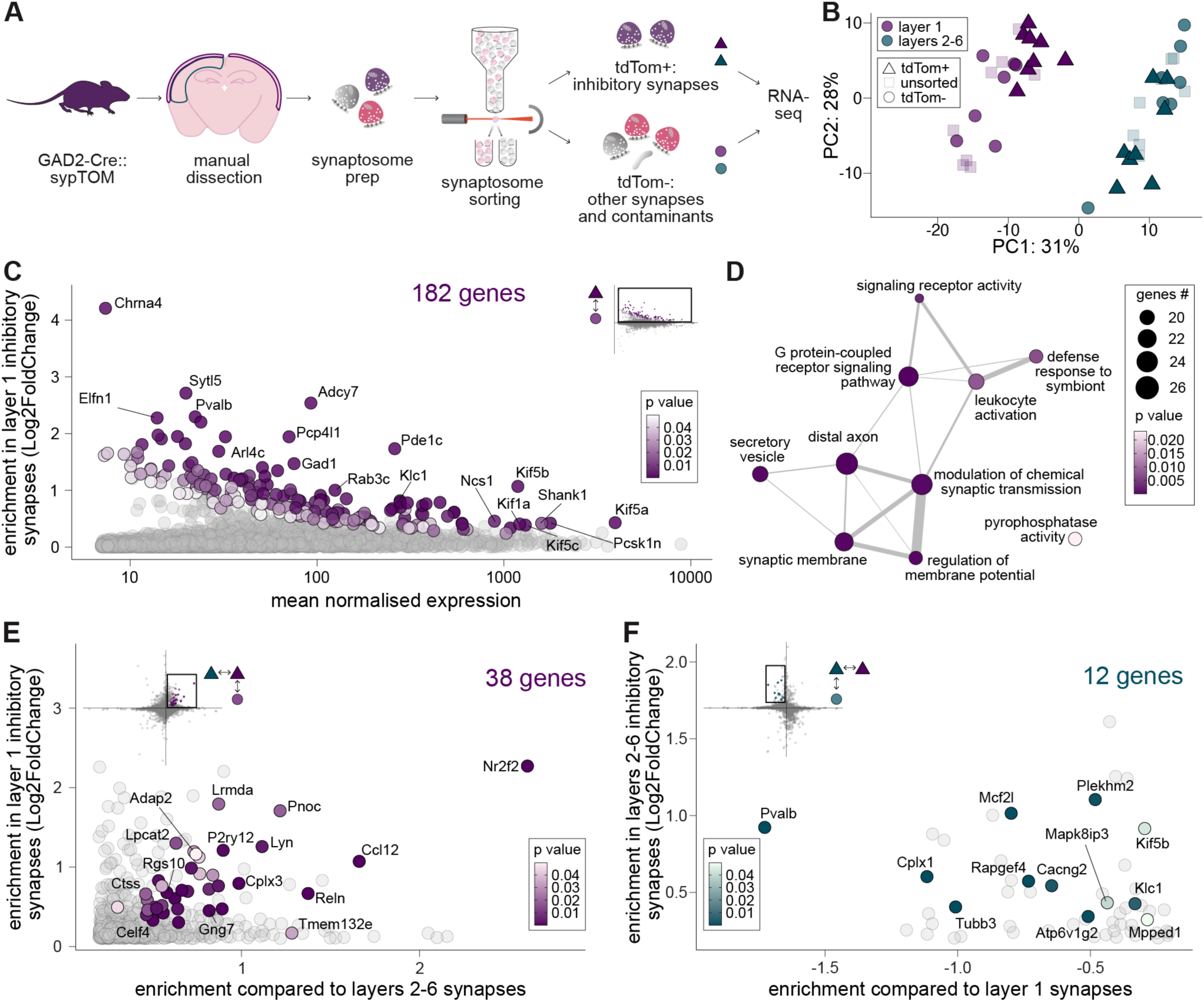
Layer 1 inhibitory synapses contain a distinct set of localized mRNAs. A) Pipeline for sorting inhibitory synapses from layer 1. B) First two components in the principal component analysis of the 500 most variable genes after correcting for individual brain variation. C) Mean normalized expression of genes against their log2FoldChange for layer 1 tdTom+ fraction (inhibitory synapses) versus its respective tdTom- (negative control): highlighted are the 182 genes that were enriched in the synapses (no log2FoldChange cut-off, adjusted p value < 0.05). D) The 10 most overrepresented GO terms associated with the genes enriched in C. The terms are ranked by gene count. Thickness of the edge represents similarity between the terms (only above 0.15). E) Scatterplot of log2FoldChange comparing the tdTom+ fraction of layer 1 and that of layers 2-6 (x axis) against the log2FoldChange in layer 1 of tdTom+ fraction against its negative control (y axis). Highlighted are the 38 transcripts that are significantly enriched in inhibitory synapses of layer 1 in both comparisons. F) Scatterplot of log2FoldChange comparing the tdTom+ fraction of layer 1 and that of layers 2-6 (x axis) against the log2FoldChange in layers 2-6 of tdTom+ fraction against its negative control (y axis). Highlighted are the 12 transcripts that are significantly enriched in layers 2-6 synapses in both comparisons.

### The synaptic protein Complexin 3 is enriched and locally synthesized in cortical layer 1

The above data can be used to discover new molecular players whose local translation could influence synaptic properties in layer 1. For example, the gene encoding the synaptic vesicle protein Complexin 3 was enriched in layer 1 in all three transcriptomic datasets (*Cplx3*; Figure 5A). Unlike Complexin 1 and 2, which are expressed at high levels in the mammalian brain, Complexin 3 is highly expressed in the retina, where it regulates exocytosis in a distinct manner ^39,40^. We thus queried whether the enrichment of *Cplx3* mRNA in layer 1 resulted in a corresponding enrichment at the protein level. Immunofluorescence of coronal sections clearly showed a higher Complexin 3 signal in layer 1 compared to the other cortical layers (Figure 5B). The difference was recapitulated when looking at Complexin 3 levels in microdissected layer 1 and layers 2-6 tissue using Western blot (Figure 5C). Next, we examined unsorted synaptosomes from the microdissected tissue, and again found that Complexin 3 was enriched in layer 1 synapses compared to those derived from layers 2-6 (Figure 5D). The enrichment of both mRNA and protein suggests that *Cplx3* is locally translated in layer 1. To directly test this, we used puromycylation proximity ligation assay (puro-PLA) in slices ^41^. We visualized abundant puro-PLA puncta, representing newly synthesized Complexin 3, in the neuropil of layer 1 (following 15 min of puromycin labelling). The signal was completely absent in slices that had not been labelled, and was drastically decreased by application of a protein synthesis inhibitor (Figure 5E). Together, these data show that Complexin 3, a synaptic protein that directly influences synaptic release, is enriched in layer 1, where it is locally synthesized, potentially at synapses.

**Figure 5.**
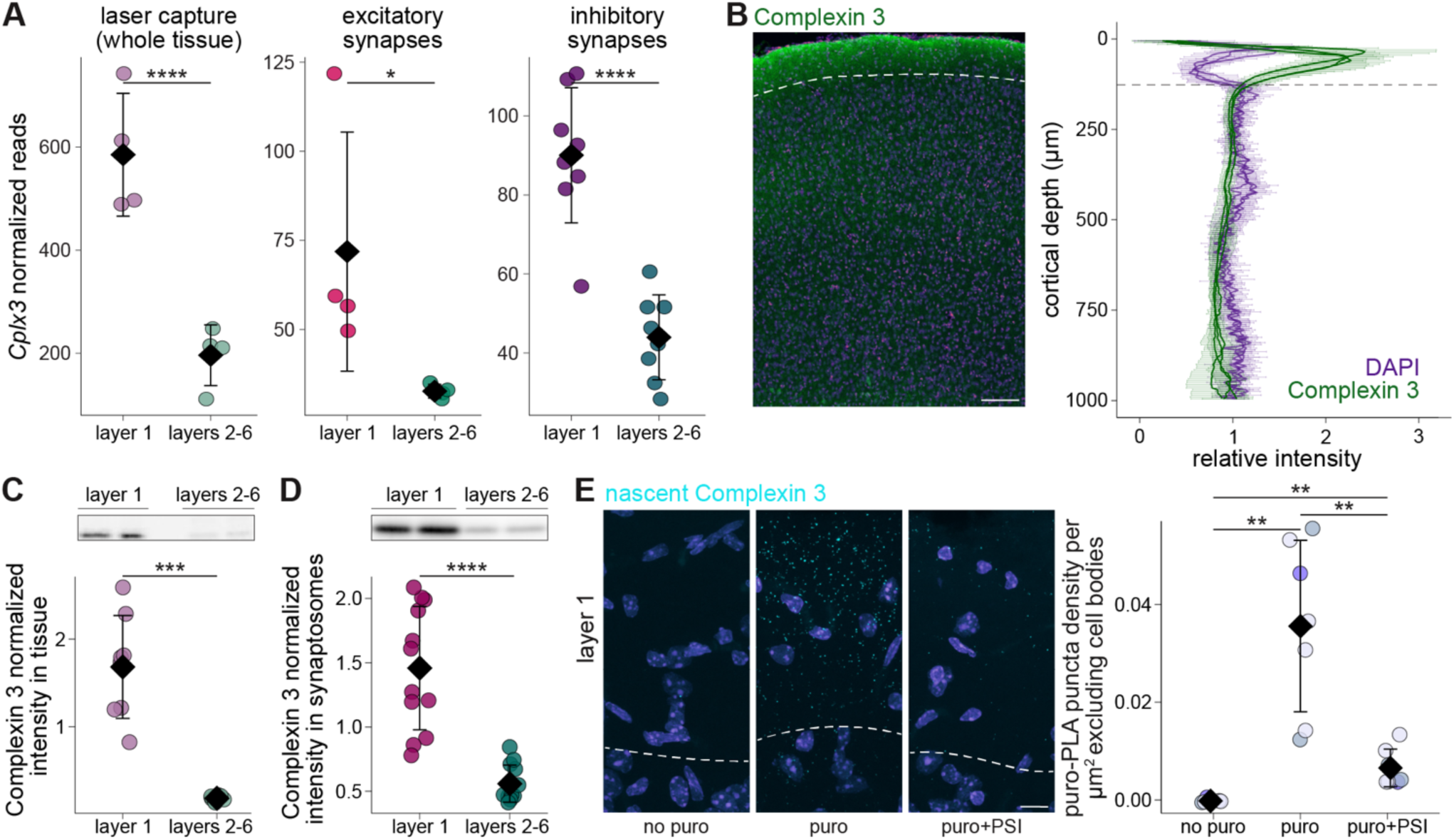
The synaptic protein Complexin 3 is enriched and locally synthesized in cortical layer 1. A) Normalized reads for the *Cplx3* gene in the three datasets of this study (n = 4 or 8 brains). Significance was calculated in the DE analysis. B) Representative image (left) and quantification (right) of Complexin 3 immunofluorescence over cortical depth. Dashed lines indicate the layer 1-2 boundary (where the nuclear density changes). Intensity measurements were averaged over 5 μm bins and are represented for each brain separately (n = 3 brain) as mean±standard deviation. Scale bar = 100 μm. C) Representative Western blot (top) and quantification (bottom) of Complexin 3 in dissected layer 1 and layers 2-6 tissue (n = 8 brains, Welch Two Sample t-test). D) Representative Western blot (top) and quantification (bottom) of Complexin 3 in synaptosomes obtained from dissected layer 1 and layers 2-6 tissue (n = 12 brains, Wilcoxon rank sum exact test). E) Representative images (left) and quantification (right) of Complexin 3 puro-PLA in cortical layer 1. Slices were treated with puromycin (1.8 μM; 15 min) +/- the protein synthesis inhibitor emetine (180 μM). Puncta density was calculated after digital exclusion of cell bodies (n = 6 or 7 slices from 3 brains, Kruskal-Wallis rank sum test followed by Wilcoxon rank sum exact test). Points represent individual slices; slices from the same mouse are represented in the same color. * p < 0.05; ** p < 0.01; *** p < 0.001; **** p < 0.0001. Points represent individual observations; the black diamond and the error bars represent the mean and standard deviation. For statistical details, see Supplementary Table 1.

### Shared transcriptomic profiles of cortical layer 1 and hippocampal stratum lacunosum moleculare

Next, we asked how the transcriptome of cortical layer 1 compared to those of the different hippocampal strata, most of which are neuropil-rich regions (Figure 6A). First, we focused on the whole tissue, comparing our laser capture data (Figure 2) with the data we recently published from microdissected CA1 strata^29^. We computed the correlation between the enrichment in layer 1 (against layers 2-6) and the enrichment in each stratum (against the whole CA1 region), discovering weak correlations between layer 1 enrichment and stratum oriens and stratum radiatum (SO and SR), a moderate negative correlation with the cell body layer, stratum pyramidale (SP), and a strong positive correlation with stratum lacunosum moleculare (SLM), the most distal hippocampal stratum (Figure 6B). We then examined significantly enriched and de-enriched genes in each group, and found a large overlap of genes regulated in the same direction in layer 1 and SLM (1996 upregulated and 1518 downregulated shared transcripts; Figure 6C). Conversely, SP had the largest set of genes regulated in the opposite direction, with 1323 downregulated transcripts being upregulated in layer 1, and 761 showing the opposing trend (Figure 6C). Which genes are enriched in both cortical layer 1 and SLM? The top GO terms associated with them were “angiogenesis”, “extracellular matrix” and “immune effector process”, suggesting that the distal location, proximity to blood vessels and exposure to immune players contributed to the transcriptomic similarity (Figure 6D). However, the large number of shared genes indicates that the similarity is not driven by a single cell-type. Indeed, the set of shared upregulated genes was composed by genes of diverse cellular origin, and only a small fraction corresponded to transcripts upregulated in brain endothelial cells, astrocytes or glia limitans^42–45^ – a cell-type found at the edge of layer 1 and SLM (Figure 6E). To test whether the similarities extend to the transcripts localized at synapses, we compared our data from layer 1 excitatory and inhibitory synapses (Figures 3E-H and 4E-F) to the synapses from each CA1 stratum^29^ (Figure S6A). The enrichment in layer 1 (against layers 2-6) of both excitatory and inhibitory synapses negatively correlated with that of SP synapses, while the highest positive correlation was found with SLM synapses, recapitulating the tissue data (Figure S6B). However, the number of commonly regulated genes was low, suggesting synaptic transcriptomes are region-specific (Figure S6C). Nonetheless, SLM synapses shared the highest number of commonly regulated transcripts (Figure S6C). Among the transcripts that were enriched in synapses of both distal layers, we found *Camk2a*, *Camk2n1*, *Homer2* and *Reln*, while those that were de-enriched in both regions included *Calm2*, ATPase subunits and voltage-gated sodium channel subunits (Figure S6D). These transcripts might be commonly regulated across brain regions to reach (or avoid) very distal synaptic sites.

**Figure 6.**
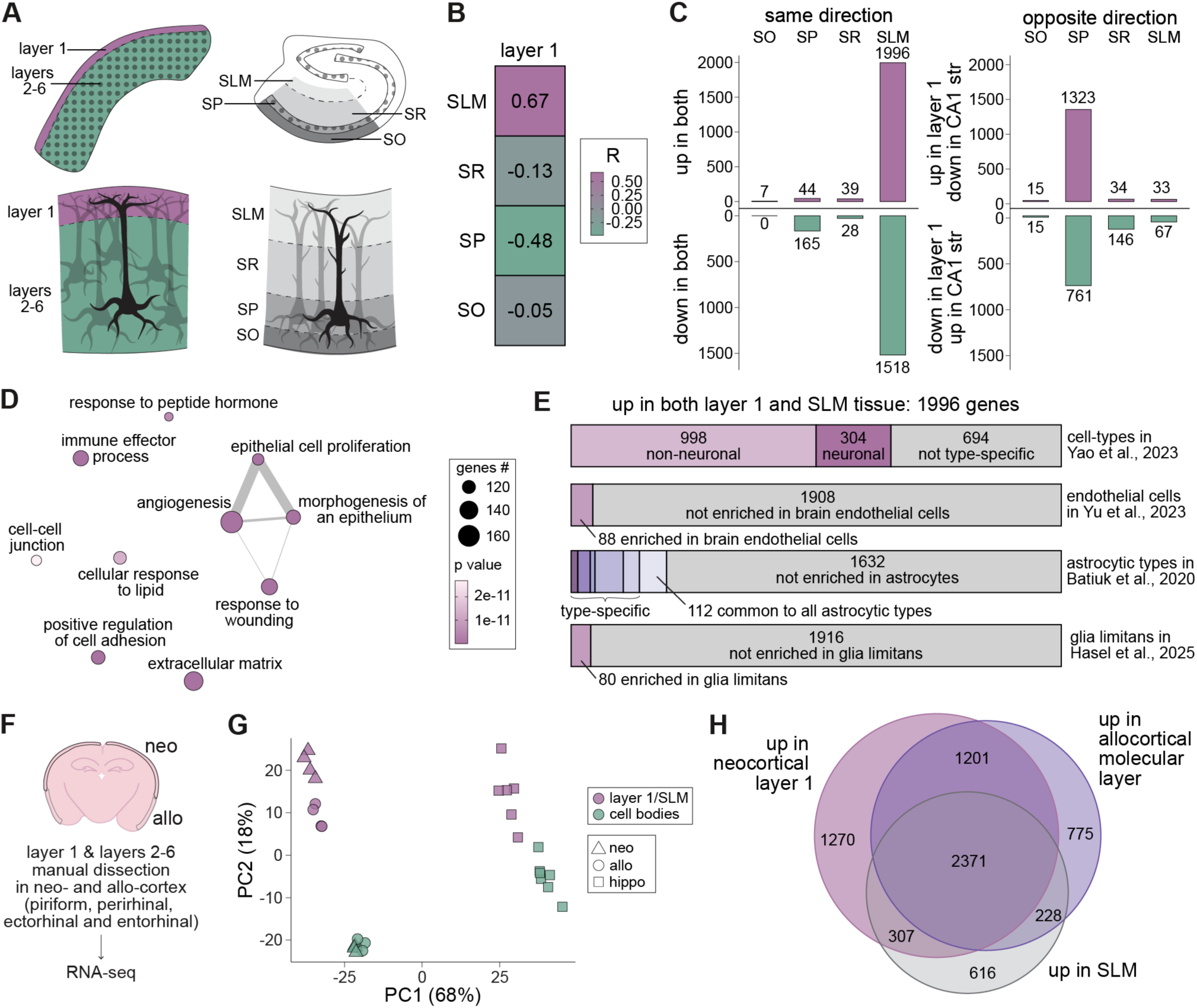
Shared transcriptomic profiles of cortical layer 1 and hippocampal stratum lacunosum moleculare. A) Drawing of hippocampus and cortex highlighting the cell bodies layers and the neuropil ones (top) and close-up of cellular organization across layers or strata (bottom). The orientation of the hippocampus is flipped vertically. B) Pearson correlation coefficients between the cortical enrichment in layer 1 in the laser capture data and the enrichment in each CA1 stratum. C) Number of genes regulated in the same (left) or opposite direction (right) between layer 1 and each CA1 stratum. D) The 10 most overrepresented GO terms associated with the genes enriched in layer 1 and SLM. The terms are ranked by gene count. Thickness of the edge represents similarity between the terms (only above 0.15). E) Overlap of enriched genes in layer 1 and SLM with genes enriched in neuronal and non-neuronal cells^43^, in brain endothelial cells^44^, in various astrocytic types^42^ and in glia limitans^45^. F) Schematic of neocortical layer 1 and allocortical molecular layer manual microdissection. G) First two components in the principal component analysis of the 500 most variable genes on the manually dissected neocortex and allocortex, merged with the CA1 SLM and SP strata. H) Overlap of enriched genes in neocortical layer 1, allocortical molecular layer and SLM compared to their respective somatic layers.

Overall, these results suggest that the transcriptomic similarity between cortical layer 1 and hippocampal SLM is neither driven by specific cell-types, nor by synaptic genes. If a specific molecular environment, involving both neuronal and non-neuronal elements, was conserved in these distal layers as the brain evolved, we expect to find it in the molecular layer of allocortical areas, such as the entorhinal, perirhinal and piriform cortex. We thus manually dissected the outermost layer of these areas from acute slices, collecting as control the respective cell body layers as well as neocortical layer 1 and layers 2-6, and processed the samples for RNA-seq (Figure 6F, S6E). The neocortical data correlated well with our previously acquired laser captured data, as expected (Figure S6F). A PCA including these data and the data from hippocampal SLM and SP revealed that, while the majority of variation was found between hippocampal and cortical samples (which were also collected and processed at different times), the data also clustered by layer, with neocortical layer 1, allocortical molecular layer and SLM separating in the same direction from the respective cell bodies (Figure 6G). Indeed, a large overlap (2371 transcripts) was found between the genes upregulated in neocortical layer 1, allocortical molecular layer and SLM (Figure 6H).

## Discussion

Local protein synthesis in the mammalian brain has primarily been studied in the hippocampus and in dissociated neuronal cultures, because of relative ease in separating the somatic and neuropil compartments. In the adult cortex, studies have been limited to synaptosomes prepared from the entire cortex^32,33^ – inevitably combining a variety of synapses from multiple areas and layers. These studies provided a general picture of cortical local protein synthesis, but no insight into how regional or synaptic specialization may occur. Here, we establish for the first time that local protein synthesis is a feature of cortical layer 1, and describe which transcripts can be locally translated to support layer 1 function, differentiating between excitatory and inhibitory synapses. By comparing the layer 1 data to their counterparts in deeper, somatic layers, we further uncover differences that could be relevant to encode layer-specific features as well as synaptic plasticity.

### The synaptic transcriptomes of layer 1

We characterized the transcriptomes of layer 1 excitatory and inhibitory synapses. Strikingly, we found more than 1000 transcripts localized in layer 1 excitatory synapses, many of which encode synaptic proteins. This suggests that local protein synthesis might contribute to the maintenance and the regulation of synaptic transmission in these synapses, which are often located far from their cell bodies. Most of these transcripts were not differentially regulated between synapses in layer 1 and those in deeper layers. However, we found that a group of synaptic genes - encoding cytosolic scaffolding proteins, calcium regulators and signaling proteins - was enriched in layer 1 synapses. These could be particularly important players for synaptic transmission in the apical tuft and in distally-projecting axons, potentially downstream of neuromodulatory inputs which are abundant in layer 1^4^. Another set of transcripts enriched in layer 1 excitatory synapses were those encoding ribosomal proteins, consistent with previous findings in other systems^33,46–48^, which could reflect a need for distal, layer 1 synapses to locally remodel their translation machinery^49^.

On the other hand, we found around 200 transcripts enriched in layer 1 inhibitory synapses, among which *Chrna4*, encoding the alpha 4 subunit of the nicotinic receptor. Local translation of *Chrna4* could regulate the synaptic sensitivity to nicotinic transmission, which in layer 1 has been shown to gate a disinhibitory circuit important for memory formation^50^. Furthermore, we found the transcripts encoding transport proteins – which regulate GABA receptors and inhibitory scaffolding proteins trafficking^51^– and those of the complement system, which might be involved in synaptic pruning^52,53^. Finally, upregulated in layer 1 inhibitory synapses compared to the deeper layers we found the transcript encoding Reelin. Reelin is expressed by layer 1 interneurons^54^ and, in the adult brain, it strongly modulates synaptic function and plasticity^55^. The presence of its mRNA at layer 1 inhibitory synapses suggests that its synthesis might be locally tuned, perhaps in response to activity.

This abundance of synaptic transcripts in layer 1 suggests that local protein synthesis could support synaptic function both under “baseline” conditions and during learning. Given the remoteness of layer 1 synapses, a local source of new proteins might be required to meet the turnover needs of some synaptic proteins. Additionally, during learning, the layer 1 circuitry needs to be reconfigured, for example with inhibitory SST+ axonal boutons being quickly eliminated^10^ while excitatory axons projecting from distal areas being potentiated^13,56,57^. Local protein synthesis might be needed for this quick rearrangement: newly synthesized calcium-dependent proteins and second messengers could mediate the potentiation of individual synaptic responses, while inhibitory synapses might be dismantled by a new pool of transport proteins.

### A shared transcriptome for distal molecular layers

By comparing our data with the transcriptome from the CA1 strata, we found a striking similarity between layer 1 and SLM, which appears conserved in the molecular layer of the allocortex. In fact, the similarities between layer 1 and SLM extend through many domains. First, they are both inhabited by apical tuft dendrites of pyramidal cells and are innervated by inputs from many distal regions^4,58^. Second, the two distal layers may rely on common synaptic circuit motifs. For example, inhibition is provided by both neurogliaform cells, with their axons densely branching locally within each layer^59,60^, and by SST+ axons arising from other layers (Martinotti and O-LM cells, respectively)^61,62^. Third, being at the pia and at the hippocampal fissure, both layers are characterized by a particular class of glial cells^42^. Finally, a Reelin extracellular gradient, high in layer 1 and SLM, is present in both regions and is essential for the localization of ion channel to the distal apical tuft of both regions^63,64^. It is thus not surprising that the transcriptomic similarity is not driven by a single cell-type or by synaptic genes. Potentially, it is the result of a conserved molecular program, involving both neuronal and non-neuronal components, that evolved to ensure the correct function of the distal, neuropil-rich molecular layer in an ancestral three-layered cortex, from which both the hippocampus and the neocortex are thought to have evolved^65^. Being far from their cell-bodies, processes in this distal layer might have been strongly influenced by the local, non-neuronal components – in line with the findings that the extracellular matrix and the immune system can regulate synaptic plasticity and neural activity in layer 1^19,66^.

### Limitations of the study

To obtain the synaptic transcriptomes of layer 1, we combined layer 1 manual dissection with fluorescence-activated synaptosome sorting. The manual dissections are likely to contain minor parts of cortical layers 2-3. However, considering the high density of synapses in layer 1, the final samples are likely composed mainly of layer 1 synapses. Furthermore, there will be a bias for presynaptic boutons, inherited from the synaptosome preparation protocol and the sorting pipeline, which relies on the expression of a presynaptic marker, although we have documented that a majority of particles also possess postsynaptic elements^37^. Therefore, the levels of postsynaptic transcripts that we detect are likely an underestimation.

Our analyses of the synaptic datasets rely on differential expression between a specific synaptic type and its negative control (tdTom+ and tdTom- fractions). The reported list of enriched genes is therefore heavily influenced by the negative control and would not include those transcripts which, despite being found at the synaptic type investigated, were also present to the same extent or more in the control. This is particularly relevant for the inhibitory synapses, as their control mostly comprised excitatory synapses (the unsorted synaptosome preparation): genes that are present in both synaptic types in similar levels will therefore not be listed in the inhibitory synaptic data. As such, our data likely represent a conservative report of these transcriptomes. Additionally, despite isolating excitatory and inhibitory synapses from a single layer, our samples are still composed of many synaptic types combined. Our dissected layer 1 tissue spans several cortical areas, and excitatory (or inhibitory) synapses will arise from a variety of presynaptic sources (for example: local vs distal, thalamic vs cortical) and postsynaptic sites (L2/3 or L5 apical dendrites, layer 1 interneurons, dendrites from deeper layers’ interneurons). This diversity might increase the variance of the data and prevent us from detecting more subtle trends and differences.

## Materials and Methods

### Animals

All procedures involving animal treatment and care were performed in conformity with the institutional guidelines, in compliance with national and international laws and policies (DIRECTIVE 2010/63/EU; German animal welfare law; FELASA guidelines; Regierungspräsidium Darmstadt approval numbers F126/2000 and F126/2002, or annex 2 of § 2 Abs. 2 Tierschutz-Versuchstier-Verordnung), and animal numbers were reported to the local authorities (Regierungspräsidium Darmstadt). All animals were housed in a 12 hour light/dark cycle and provided with food and water *ad libitum*. For synaptosome sorting experiments, homozygous mice from the conditional SypTOM mouse line (RRID:IMSR_JAX:012570) were crossed with homozygous NEX-Cre^67^ (obtained from the Nave lab at the MPI for multidisciplinary sciences in Göttingen) or GAD2-Cre^68^ (RRID:IMSR_JAX:010802) mice to obtain NEX-Cre::SypTOM or GAD2-Cre::SypTOM mice heterozygous in both loci. NEX-Cre::3xMetRS* mice were obtained as described in Alvarez-Pardo et al.^35^.

### Perfusions and sectioning

Wild-type C57BL/6J mice between 9 and 16 weeks old were terminally anesthetized with an i.p. injection of 300 mg/kg ketamine and 20 mg/kg xylazine. After losing all reflexes, the mice were transcardially perfused first briefly with ice-cold DPBS (Thermo Fisher Scientific #14040174), then with ice-cold 4% paraformaldehyde (PFA, Thermo Fisher Scientific #28908), 4% sucrose in PBS. For some immunofluorescence (IF) experiments, the extracted brains were post-fixed in the fixative solution overnight at 4 °C, then washed twice in PBS, and sectioned to 40 or 50 μm coronal slices using a vibratome (Leica VT1200S; 0.16 mm/s). For FISH and other IF experiments, the fixed brains were extracted and further incubated in the fixative solution for 1 hour at room-temperature. After two washes in PBS, the brains were cryoprotected over two nights in 30% sucrose in PBS. All buffers were made with RNAse-free reagents. The brains were then cryosectioned to 40 μm coronal slices using a microtome (Zeiss Hyrax S50). For FISH, the slices were washed, post-fixed at room temperature for 10 minutes in a fixative solution (4% PFA, 5.4% glucose, and 0.01 M sodium metaperiodate in lysine-phosphate buffer) and immediately used.

### Immunofluorescence

Slices were permeabilized for 3 to 5 hours in 0.1% triton in blocking buffer (4% goat serum in PBS), and then incubated in primary antibody solution 24-72 hours at 4 °C with shaking (the following antibodies diluted 1:1000 in blocking buffer: rabbit anti-NeuN from abcam #ab177487; chicken anti-GFAP from Novus #NBP1-05198; mouse anti- S100β from abcam #ab11178; rabbit anti-Complexin 3 from Proteintech #16949-1-AP). The slices were then washed for 10 minutes in PBS three times and incubated in secondary antibody solution overnight at 4 °C (the following antibodies diluted 1:1000 in blocking buffer: goat anti-rabbit-Alexa488 #A11008; goat anti-chicken-Alexa488 #A11039; goat anti-mouse-Alexa488 #A11001; goat anti-chicken-Alexa647 #A21449; all acquired from Thermo Fisher Scientific). The slices were washed for 10 minutes in PBS twice, incubated in DAPI (Thermo Fisher Scientific #D1306) diluted in PBS for 10 minutes, washed again twice for 10 minutes in PBS and then mounted in Aqua-PolyMount (Polysciences #18606-20) onto SuperfrostTM Plus Adhesion slides (Epredia #J1800AMNZ) covered by glass coverslips (VWR #631-0138).

### Fluorescence *in situ* hybridization (FISH)

FISH was performed using the QuantiGene ViewRNA kit from Thermo Fisher Scientific (#QVP0011): slices were permeabilized for 20 minutes using the kit’s detergent buffer, and detection probes were incubated overnight at 40 °C (ViewRNA ISH probe sets: VC1-11005, VF1-11494, VB6-3200449, VC1-18640, VB1-15336, VB1-3048317, VB1-10350, VC6-10835). Preamplification, amplification, and label probes were applied according to the manufacturer’s recommendations, with three washes of 5 minutes between each step using the provided wash buffer. All reagents used until this step were RNAse-free. Finally, the slices were washed in PBS, stained with DAPI for 20 minutes at room temperature, washed again in PBS and mounted as described above.

### Acute slice preparation

Wild-type C57BL/6J, NEX-Cre::3xMetRS*, NEX-Cre::SypTOM or GAD2-Cre::SypTOM mice (5-10 weeks old) were anesthetized with isoflurane and quickly decapitated. The brains were dissected in ice-cold sucrose ACSF (87 mM NaCl, 25 mM NaHCO_3_, 1.25 mM NaH_2_PO_4_, 2.50 KCl, 10 mM glucose, 75 mM sucrose, 0.5 mM CaCl_2_, 7 mM MgCl_2_, continuously bubbled with 95% O2 and 5% CO_2_) and cut to 300 μm coronal slices with a vibratome (Leica VT1200S; 0.08 mm/s with 1.0 mm vibration). Slices between AP -0.5 and AP -3.5 from bregma were selected. For metabolic labelling experiments, slices recovered for at least 1 hour in room-temperature ACSF (125 mM NaCl, 25 mM NaHCO_3_, 1.25 mM NaH_2_PO_4_, 2.50 KCl, 10 mM glucose, 2 mM CaCl_2_, 1 mM MgCl_2_, continuously bubbled with 95% O2 and 5% CO_2_). For synaptosome sorting experiments, slices recovered at 36°C for 30 minutes in sucrose ACSF before being transferred to ACSF.

### Layer 1 manual microdissection

For all manual microdissection experiments, the slices were incubated in recovery solution (sucrose ACSF or ACSF) containing NucBlue Live ReadyProbes Reagent (Thermo Fisher Scientific #R37605, 1.5-2.5 drops/10 mL) for at least 30 minutes. The slices were then transferred onto a clear dissection Petri Dish (Scintica Instrumentation Inc. #DD-90-S-3PK) and layer 1 was dissected using a clean microsurgical knife (WPI #500250) under an Axiocam 506 mono (Zeiss) which enabled the visualization of the dye marking the cell nuclei. The border between layer 1 and layer 2 was recognizable because of the change in cell body density. Layer 1 was dissected in neocortical areas, starting from the rhinal fissure up until the interhemispheric fissure in both hemispheres. The corresponding layers 2-6 were collected from a subset of slices. For metabolic labelling experiments, the dissected tissue was kept in ACSF at room temperature until layer 1 was dissected from all slices. For synaptosome experiments, the tissue was directly collected into gradient medium (GM; 0.25 M sucrose, 5 mM Tris-HCl, 0.1 mM EDTA, supplemented with protease inhibitor from Millipore #539134 - diluted 1:750 - and SUPERase•InTM RNase Inhibitor from Thermo Fisher Scientific #AM2696 - diluted 1:80). For the manual dissection of the allocortex, the outermost layer was separately collected in the regions of perirhinal, entorhinal, ectorhinal and piriform cortex that were accessible. The corresponding somatic layers were collected in parallel.

### Metabolic labelling in slices or dissected tissue

Dissected layer 1 and layers 2-6 were incubated for 15 minutes in ACSF containing 18 μM (10 μg/mL) puromycin (Sigma-Aldrich #P9620) at 36°C, or for 1 hour in ACSF containing 1 mM ANL (Iris Biotech #HAA1625) at 34°C. Then the tissue was transferred to ACSF and briefly centrifuged, before the ACSF was removed and the tissue snap-frozen. For the protein synthesis inhibitor control, slices were transferred to ACSF containing either 180 μM (100 μg/mL) emetine (for puromycin experiments, Cayman Chemical #Cay21048-50) or 355 μM (100 μg/mL) cycloheximide (for ANL experiments, Sigma-Aldrich #C7698) before microdissection, and kept in the same solution during labelling. For puromycin experiments, frozen tissue was lysed in RIPA buffer (Thermo Fisher Scientific #89901) supplemented with protease and phosphatase inhibitors (1:100, Thermo Fisher Scientific #78442) and Benzonase (1:1000, Sigma-Aldrich #E1014) by sonication and used directly for SDS-PAGE. For ANL experiments, the frozen tissue was homogenized in PBS (pH 7.4) containing 1% SDS, 1% Triton X-100, Benzonase (1:1000) and protease inhibitors (EDTA free, 1:4000) with a hand-held mini homogenizer, heated to 75 °C for 15 min and centrifuged at 17,000 x *g* for 15 min at 10 °C. The supernatant was collected, protein concentration measured and equal amounts of protein were used for the alkylation and click reaction^69^. For puro-PLA experiments, full slices (without dissection) were transferred to ACSF containing 1.8 μM (1 μg/mL) puromycin for 15 minutes, then transferred to ACSF for another 10 minutes before being fixed in a solution containing 4% paraformaldehyde, 4% sucrose in PBS overnight. A subset of slices was incubated in ACSF containing 180 μM (100 μg/mL) emetine for 1 hour before labelling; another subset was fixed without any labelling.

### Alkylation and click reaction

The ANL-treated samples were diluted 1:2 (v/v) in PBS (pH 7.8) with protease inhibitors (1:2000) and iodoacetamide (Sigma-Aldrich #I6125) was added twice to a final concentration of 20 mM, incubating the samples for 2 hours in the dark at room temperature with rotation after each addition. The buffer was then exchanged to click chemistry buffer (PBS pH 7.8, 0.04% SDS, 0.08% Triton X-100, protease inhibitors 1:4000) using PD SpinTrap G-25 columns (Cytiva #28918004). Each click reaction was set up with 120 μL of sample by adding the following reagents in order and vortexing for 20 seconds after each addition: 1.5 μL of triazole ligand (stock 200 mM, Sigma-Aldrich #678937), 1.5 μL of FLAG alkyne (stock 50 mM, [Proparagylglycin]DYKDDDDKDYKDDDDK[COOH], custom made from Thermo Fisher Scientific), and 1 μL of Cu(I) bromide (stock 10 mg/mL, Sigma-Aldrich #254185, freshly dissolved in DMSO, Sigma-Aldrich #276855). Once everything was added, the reactions were kept rotating in the dark overnight at room temperature. The samples were then centrifuged for 5 minutes at 17,000 x *g* and the supernatants were kept. For layer 1, two reactions were set up for each sample, then pooled together and concentrated in a SpeedVac vacuum concentrator for 30-45 minutes at 45 °C. The samples were then processed for SDS-PAGE.

### Synaptosome preparation

The dissected tissue was collected in GM and then transferred to a clean 1.0 mL glass WHEATON® Dounce Tissue Grinder and homogenized with 20 strokes with the “loose” and 20 with the “tight” pestle. The samples were then centrifuged for 10 minutes at 4 °C at 1,000 x *g*. The S1 supernatant was layered onto a density gradient (23%, 10% and 3% Percoll in GM buffer). The gradient was centrifuged in an Avanti J-26S XPI centrifuge with a JA-25.50 rotor (Beckman Coulter) for 5 minutes at 4 °C at 32,500 x *g* with maximum acceleration and minimum deceleration. The F2/3 fraction, the third from the top, was collected with a syringe and a blunt cannula, and used for metabolic labelling or as input for the Fluorescence-Activated Synaptosome Sorting (FASS). In some experiments, an aliquot of the original homogenate, of the S1 supernatant and of the F2/3 fraction were kept for SDS-PAGE.

### Metabolic labelling in synaptosomes

Layer 1 and layers 2-6 synaptosomes were washed twice in PBS by centrifugation (10 minutes at 16,000 x *g*), resuspended in PBS and divided into two 30 μL reactions (+/- emetine) containing 2 mM ATP (from a 50 mM stock made fresh every two weeks), 125 mM HEPES, 10 mM MgCl2, 20 μM amino acids (Promega #L4461), 240 μM puromycin (from a 3 mM stock), 4 mM emetine (only in half of the reactions) in PBS. Protein synthesis was labelled at 35 °C for 15 minutes with gentle shaking (350 rpm). The synaptosomes were then lysed in 1% SDS, NuPAGE LDS sample buffer and sample reducing agent (Invitrogen #NP0007 and #NP0009) were added, the samples were evaporated for 5 minutes at 95 °C with gentle shaking (350 rpm) and immediately used for SDS-PAGE.

### SDS-PAGE and Western blot

If needed, sample protein concentration was measured using the Precision Red advanced protein assay (Cytoskeleton #ADV02) or Pierce BCA Protein Assay (Thermo Fisher Scientific #23225). The same amounts of proteins were mixed with NuPAGE LDS sample buffer and sample reducing agent (Invitrogen #NP0007 and #NP0009). After incubation for 5 minutes at 85-95 °C, the samples were separated onto Novex 4-12% Tris-Glycine WedgeWell gels or BOLT 12% Bis-Tris Plus WedgeWell gels (Thermo Fisher Scientific #XP04205BOX, #XP04200BOX, #NW00127BOX) by running for 1-2 hours at 100-150V in Tris-Glycine SDS running buffer (25 mM Tris, 192 mM glycine, and 0.1% SDS) or NuPage MOPS SDS running buffer (Thermo Fisher Scientific #NP0001). Proteins were then transferred onto nitrocellulose membranes using the Bio-Rad Trans-Blot Turbo system and its transfer packs (Bio-Rad #1704158 or #1704159). Total protein was visualized using the Revert 700 Total Protein stain (Licor #926-11021) or Ponceau staining (Cell Signaling technology #59803S) according to manufacturers’ instructions. After de-staining, the membranes were blocked with Intercept PBS blocking buffer (Li-Cor Bio, 927-70001) for 1 hour at room temperature and incubated overnight at 4 °C in primary antibodies (diluted in blocking buffer: rabbit anti-puromycin from abcam #ab315887, 1:1000; rabbit anti-FLAG from Cell Signaling Technologies #CS2368S, 1:1000; rabbit anti-NeuN from abcam #ab177487, 1:1000; rabbit anti-Histone H3 from abcam #ab1791, 1:1000; mouse anti-Synaptophysin from Sigma-Aldrich #S5768, 1:2500; mouse anti-PSD95 from Thermo Fisher Scientific #MA1-046, 1:1000; rabbit anti-Complexin 3 from Proteintech #16949-1-AP, 1:1000). After three washes in TBS containing 0.1% triton, the membranes were then incubated in secondary antibodies (LICOR antibodies diluted 1:5000: goat anti- mouse-IRDye680 #926-68020, goat anti-rabbit-IRDye680 #926-68071, goat anti- mouse-IRDye800 #926-32210, goat anti-rabbit-IRDye800 #926-32211; Cell Signaling antibodies diluted 1:3000: horse anti-mouse-HRP #CS7076, goat anti- rabbit-HRP #CS7074) diluted in Intercept PBS blocking buffer or in 5% milk in TBS for 1 hour at room temperature. After three washes in TBS containing 0.1% triton, the membranes were either imaged with the Li-Cor Odyssey 9120 gel imaging system, or developed using the SuperSignal West Femto (Invitrogen, 34094) and imaged using the Azure 280 imaging system. Quantification was done using the “Gels” functions in ImageJ^70^ and was plotted and analyzed in R^78^. All intensity values – signal and total protein – were first normalized by the average intensity of each gel (which included all conditions); then the normalized signal was divided by the normalized total protein.

### Laser capture microdissection

Wild-type C57BL/6J mice between 9 and 16 weeks old were anesthetized with isoflurane, and their brains quickly dissected. After a brief immersion in RNALater (Invitrogen #AM7024), the brains were positioned in a plastic mold, covered with cryo- embedding compound (Ted Pella Inc #27300) and flash-frozen in isopentane which had been previously cooled between -60 °C and -70 °C on dry ice. After two minutes in isopentane the brains were transferred and stored at -76 °C until cryosectioning. 10 μm sections were cut using a cryostat (CM3050, Leica), mounted on individual Menzel-Gläser Superfrost Ultra Plus slides (Thermo Fisher Scientific) and immediately processed for laser capture or stored at -76 °C until used (maximum 48 hours after cryosectioning). Right before laser capture, each section was fixed by immersion in 75%, 95%, and 100% ethanol (1 minute each) and rehydrated by immersion in 95%, 75% ethanol and PBS (1 min each). Laser capture was performed at a PALM MicroBeam Microscope (Zeiss), and the energy and focus of the laser beam were adjusted before each experiment with a dummy section to ensure efficient laser capture of the tissue. The entire layer 1 from each section was first laser captured in an adhesive cap (AdhesiveCap 200 clear, Zeiss), collecting only the outermost ∼50 μm. 1 μL of lysis buffer (from the Absolutely RNA Nano Kit, Agilent Technologies; #400753), pre-mixed with β-Mercaptoethanol as per manufacturer instruction, was added to the sample which was then stored on dry ice. Layers 2-6 were then collected from the same section in a different tube (only auditory cortex and temporal association cortex). The process was repeated for 12 sections from the same brain (all between bregma AP -1.5 and -3.5).

### Laser capture library preparation

All layer 1 and layers 2-6 samples were pooled together and processed for RNA extraction, which was done using Absolutely RNA Nano Kit (Agilent Technologies; #400753) and included a DNAse I step. Total RNA was eluted in 15 μL RNase free water and its quality was assessed using RNA 6000 Pico Kit (Agilent Technologies, #5067-1513) on a Bioanalyzer (Agilent Technologies). Libraries were prepared from total RNA ranging from 1.2 to 6.6 ng using NEBNext Ultra II Directional RNA Library Prep Kit for Illumina (#E7760S) and NEBNext® Multiplex Oligos for Illumina® (96 Unique Dual Index Primer Pairs, #E6440). mRNAs were enriched using the NEBNext poly(A) mRNA Magnetic Isolation Module (#E7490). Final cDNA libraries were assessed using Agilent High Sensitivity DNA assay (Agilent Technologies, #5067-4626) and sequenced on Illumina NextSeq2000, using a paired-end 75-bp sequencing run.

### Manually dissected tissue library preparation

Manually dissected neocortical layer 1 and layers 2-6 and allocortical molecular layer and somatic layers were frozen on dry ice and stored at -80 °C. Tissue was resuspended in 1 mL TRIzol, transferred to a clean 1.0 mL glass WHEATON® Dounce Tissue Grinder, homogenized using a ‘loose’ and ‘tight’ pestle for 20 strokes each then transferred to an RNAse-free 2 mL tube and further triturated 20x using a 23 G needle. All samples were incubated for 5 minutes at room temperature and centrifuged for 3 minutes at 16,100 × *g*. RNA was purified and extracted from the supernatant using Direct-Zol RNA Miniprep Kit (ZymoResearch #R2051). RNA quantity and quality were measured on a Qubit fluorometer (Invitrogen #Q33216) and on a 2100 Bioanalyzer RNA Pico chip. Libraries were prepared from 25 ng of total RNA using NEBNext Ultra II Directional RNA Library Prep Kit for Illumina (E7760), Poly(A) mRNA Magnetic Isolation Module (E7490) and Multiplex Oligos for Illumina Set A (E6440). Final cDNA libraries were assessed as above and sequenced on Illumina NextSeq2000, using P2 reagents with a paired-end 61-bp sequencing run.

### Fluorescence-activated synaptosome sorting (FASS)

FASS was performed on a FACSAria Fusion running FACSDiva (BD Biosciences) with the following settings: 488nm laser (for FM4-64), 561nm laser (for tdTomato signal), sort precision (0-16-0), FSC701(317 V), SSC (488/10 nm, 370V), PE ‘‘tdTomato’’ (586/15 nm, 470V), PerCP ‘‘FM4-64’’702(695/40 nm, 470), thresholds (FSC = 200, FM4-64 = 700). Synaptosomes were mixed with a membrane dye (FM4-64; Thermo Fisher Scientific #T13320, diluted 1: 100 to create a stock, of which 1.5 μL were mixed with 1 mL of F2/3 solution) and then sorted through a 70 μm nozzle at ∼20,000 events/s and a flow rate of < 5 μL/min with FACSFlow as sorting buffer (BD Biosciences #12756528). They were collected on a Whatman GF/G glass microfiber filter (Cytiva #1825-090) supported by a PE drain disk (Cytiva, #231100), which had previously been punched out together using a 4 mm Militex® biopsy punch (#15110- 40). The filter was washed with RNAsecure (Thermo Fisher Scientific #AM7006, diluted 1:5) in sorting buffer prior sorting 2 million synaptosomes per condition. The filter was then washed with 1 mL of RNAse-free PBS and stored at - 76 °C until library preparation. For each tissue sample (layer 1 or layers 2-6 from each mouse, or control and astro samples from each mouse), 3 filters were collected: cell-type specific synapses were sorted based on the particle size (using the membrane dye) and the tdTomato signal (“tdTom+”); a control fraction which included other synapses and similarly-sized contaminants was sorted based on particle size and the absence of the tdTomato signal (“tdTom-”); finally, the F2/3 precursor fraction was also sorted, just based on particle size but independently of the tdTomato signal (called “unsorted”, only used for quality evaluation and PCA plots). All procedures were conducted on ice.

### Synaptosomes library preparation

First, DNA was digested and membranes were lysed by incubating each filter for 20 minutes at 20 °C in 7.5 μL of 1 mM DTT, 1 U/μL Recombinant RNAse Inhibitor (Takara Bio #2313A), 1x SingleShot Lysis Buffer and 1x DNAse Solution (Bio-rad #1725080), followed by enzyme inactivation for 5 minutes at 75°C. Individual reverse transcription primers (see list below) at a concentration of 1 mM were then hybridized in a solution of 10 mM DTT, 1mM dNTPs, 1.5 mM dCTPs and 1 U/μL Recombinant RNAse Inhibitor for 5 minutes at 65 °C, and immediately transferred on ice. Reverse transcription was carried out by SuperScript IV (10 U/μL) in 1x SuperScript RT buffer (Thermo Fisher Scientific #18090050), 6mM MgCl_2_, 1 M Betaine (Merck Millipore #B0300), and 1 μM template switch oligo (5’-/5BiosG/ AAGCAGTGGTATCAACGCAGAGTGTCGTGACTGGGAAAACCCTGGCrGrGrG-3’) for 10 minutes at 55 °C, following by enzyme inactivation for 10 minutes at 80 °C. The sample was then mixed with 1x of the DNA polymerase mix (Roche, #07958935001) and 0.1 μM PCR primer (5’-AAGCAGTGGTATCAACGCAGAGT-3’), and PCR was performed in the following steps: 3 minutes of initial incubation at 98 °C, 18 cycles of 20 seconds at 98 °C, 10 seconds at 67 °C and 6 minutes at 72°C, and at last 5 minutes of extension phase at 72 °C. Between all the steps until the PCR the samples were kept on ice. The microfiber filter was then removed by spinning down the sample in a Costar® Spin-X® filter column (Corning #8162) for 1 minute at 3,000 x *g*. DNA was size-selected using 0,6x sample volume of AMPure XP beads (Beckman Coulter #A63881) according to manufacturer’s instructions, and resuspended in 7 μL elution buffer (Qiagen #19086). DNA concentration was measured using the High Sensitivity DNA Assay (Thermo Fisher Scientific, #Q32854) on a Qubit Fluorometer (Thermo Fisher Scientific), and the quality assessed using Agilent High Sensitivity DNA assay (Agilent Technologies #5067-4626) on a Bioanalyzer (Agilent Technologies). Concentrations of indexed cDNAs were normalized to 0,1ng/µl and samples were pooled together for subsequent library preparation using Nextera XT DNA kit (Illumina, #FC-131-1096). Briefly, 5 μL of pooled DNA was tagmented by adding 10 μL of Tagment DNA buffer and 5 μL of Amplicon Tagment Mix, and incubating the mixture for 5 minutes at 55 °C. The reaction was stopped by adding 5 μL of Neutralize Tagment Buffer, then spinning down by centrifugation for 1 minute at 280 x *g* and incubating for 5 minutes at room temperature. The samples were added to the Nextera PCR mixture: a total of 50 μL per reaction included 15 μL Nextera PCR Master Mix, 0.2 μM P5-ISPCR custom primer (5’-AATGATACGGCGACCACCGAGATCTACACGCCTGTCCGCGGAAGCAGTGGTATCAACG-3’), 0.2 μM i7 adaptor with specific sequences (see list below), and 25 μL of tagmentation reaction. The reaction was carried out with the following steps: 3 minutes at 72 °C of pre-PCR incubation, 30 seconds at 95 °C for an initial denaturation, and 12 cycles of 10 seconds at 95 °C, 30 seconds at 55 °C and 30 seconds at 72 °C. Finally, samples were incubated for 5 minutes at 72 °C. The DNA was purified with AMPure XP beads, eluted with 15 μL of elution buffer (QIAGEN). Following the estimation of concentrations and average sizes, all libraries were pooled in equimolar amounts for a sequencing run. Paired-end sequencing was performed on Illumina NextSeq2000 with P2 (100) reagents and a custom sequencing primer for Read 1 (5’- GCCTGTCCGCGGAAGCAGTGGTATCAACGCAGAGTAC -3’). The following parameters were used: Read1 = 26 bp, Read2 = 82 bp, Read1 Index = 8 bp, Read2 Index = 0 bp.

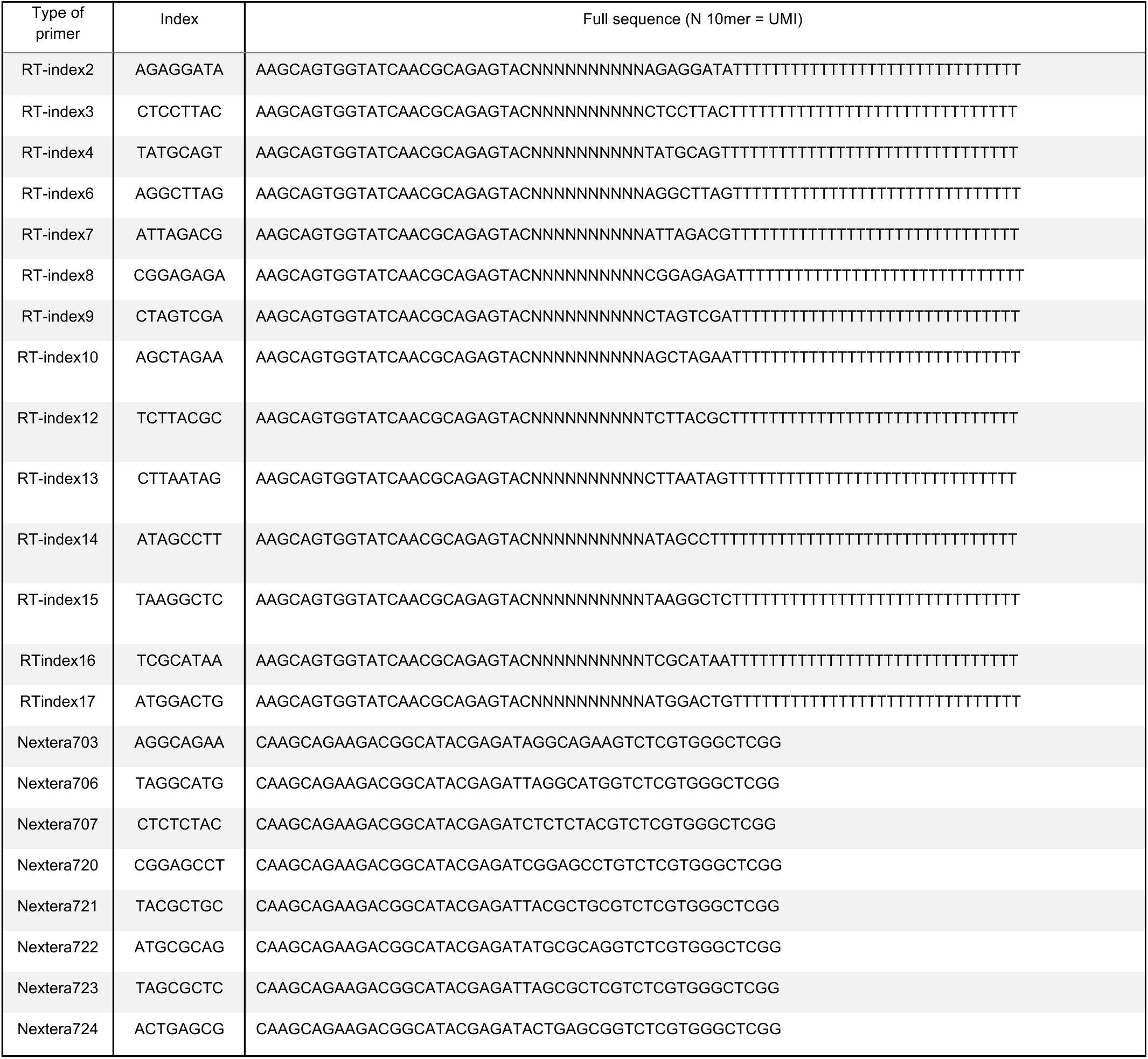

### Puro-PLA

Fixed labelled slices were washed twice in PBS and cryoprotected at least overnight in 30% sucrose in PBS. The brains were then cryosectioned to 40-50 μm coronal slices using a microtome (Zeiss Hyrax S50). Slices were then permeabilised overnight at 4°C in 0.1% triton in blocking buffer (4% goat serum in PBS), then incubated in primary antibody solution overnight at 4°C (the following antibodies diluted 1:1000 in blocking buffer: rabbit anti-puromycin from abcam #ab315887; mouse anti-Complexin 3 from Santa Cruz #sc-365941) and washed three times in PBS at room temperature. Proximity ligation was carried out using Duolink reagents (Sigma-Aldrich #DUO92007). Slices were incubated in secondary antibody solution for 2 hours at room temperature (the following antibodies diluted 1:10 in antibody buffer from the kit: anti-mousePLA-minus and anti-rabbitPLA-plus, Sigma-Aldrich #DUO92002 and #DUO92004). After five washes in buffer A, slices were incubated in the ligation reaction for 1 hour at 37°C in a humidified chamber. After three washes in buffer A, slices were then incubated in the amplification reaction for 1 hour at 37°C in a humidified chamber. Finally, slices were washed three times in buffer B, then three times in PBS, labelled with DAPI and mounted as described above.

### Confocal imaging

Immunofluorescence, FISH and puro-PLA images were acquired on an LSM880 confocal microscope (Zeiss) equipped with a 20x/0.8 air, 40x/1.3 or 63x/1.4 oil objective, or on an LSM780 confocal microscope (Zeiss) with a 10x/0.45 air objective. Laser lines and detectors’ spectral windows were selected depending on the used fluorophores. Z- stacks covering the entire slice were acquired using Zen 2.3 SP1 FP3 (black) (Zeiss, version 14.0.29.201). Imaging parameters were adjusted to prevent saturation and were kept consistent within but not between sessions. In FISH and puro-PLA experiments, pixel size was determined according to the Nyquist criterion as calculated by the ZEN software. In every session a negative control was included. For the FISH data, the negative control was used to determine which threshold for puncta detection to use in the rest of the images of that experiment. Maximum intensity projections were used for analysis and, after contrast adjustment, for display.

### Image analysis

Complexin 3 intensity and puncta quantification for FISH and puro-PLA was performed in ImageJ2 (version 2.14.0/1.54f)^70^ using in-house scripts. For puncta quantification, a mask of layer 1 was manually created based on the DAPI staining, which was also used to automatically remove the areas of the image that included cell bodies. If the automatic detection did not work, cell bodies were selected manually. The remaining area (neuropil) was used for quantification of puncta, which were identified after manual background subtraction and median filtering using the “Find Maxima…” function (with prominence 100, 250 and 500 for FISH; 4000 for puro-PLA). For Complexin 3 intensity quantification, DAPI and Complexin 3 intensities were measured across the cortical layers, normalized by the mean of the field of view, and averaged in 5 μm bins. Quantification over several fields of view from the same animal were then averaged. All data were plotted and analyzed in R.

### Preprocessing of laser captured and microdissected tissue data

Raw BCL files from Illumina sequencing were processed on a high-performance cluster (1 node, 32 CPU cores, SLURM job scheduler). The preprocessing pipeline comprised demultiplexing, read trimming, alignment, and gene expression quantification. Raw data were demultiplexed using bcl2fastq2 (version 2.20.0.422), with sample sheet specification and the --no-lane-splitting option. Output FASTQ files were organized per sample. Demultiplexed FASTQ files were quality-filtered and trimmed with fastp (version 0.24.0)^71^, using: adapter trimming: AGATCGGAAGAGCACACGTCTGAACTCCAGTCA (R1), AGATCGGAAGAGCGTCGTGTAGGGAAAGAGTGT (R2); quality-based trimming (window size: 4, min mean quality: 20), polyX trimming (min length: 10), and low- complexity filtering (threshold: 30); reads shorter than 51 bases or average quality below 20 were discarded; QC reports were generated in JSON and HTML formats. Clean reads were aligned to the Mus musculus mm39 genome using STAR (version 2.7.10a)^72^. BAM files were sorted by coordinates and indexed using samtools (version 1.21)^73^. Aligned BAM files were processed with featureCounts (version 2.0.3)^74^ with the following parameters: exon-level counting (-t exon), gene_id as the feature, high-quality read filtering (-Q 255), paired-end mode (-p). Output was a sample-wise gene count matrix. Pipeline steps were automated via a configuration file provided at runtime. Intermediate and final outputs were organized into structured directories. The preprocessing pipeline ensured high-quality, reproducible data for downstream analysis.

### Preprocessing of sorted synaptosome data

Barcoded synaptosome RNA-sequencing data were processed using an automated pipeline incorporating demultiplexing, quality filtering, and alignment^75^. Raw sequencing data were demultiplexed using ‘bcl2fastq’, converting base- call files into FASTQ format. Reads were filtered with ‘fastp’^71^ to remove adapters, low-complexity sequences, and low-quality reads. Alignment and quantification were performed using ‘STAR’ (v2.7.11b)^72^ in ‘STARsolò mode with ‘--soloType CB_UMI_Simplè, integrating read mapping, barcode error correction (‘--soloCBmatchWLtype 1MM’), UMI deduplication (‘--soloUMIdedup 1MM_All’), and gene expression quantification (‘--soloFeatures GeneFull_Ex50pAS’). The data were aligned against the *Mus musculus* (mm39) reference genome (Genome Reference Consortium) and annotation from UCSC Genome Browser. The resulting BAM files, gene expression matrices, and alignment logs were systematically organized for downstream analyses. The code to the pipeline is publicly available under: https://gitlab.mpcdf.mpg.de/mpibr/schu/synaptosort.

### Analysis of sequencing data

Once raw counts were obtained for each sample, analysis of the data was done using the DESeq2^76^ and clusterProfiler^77^ packages in R^78^, and plotting with the ggplot2^79^ and enrichplot^80^ packages. Transcripts per million were calculated by first obtaining reads per kilobase (normalizing by gene length), then normalizing by the total number of reads in the library. Normalized counts values were obtained using the respective function in DESeq2. In the laser capture dataset, genes were considered detected in a layer if their average normalized counts in those samples was above 4. In the synaptosome data, genes whose average was below 4 normalized counts in all conditions were excluded *a priori*. PCA plots were made using the DESeq2 function ‘plotPCÀ. To correct for individual brain variation, the function ‘removeBatchEffect’ from the limma package^81^ was used. Synaptic gene lists were obtained from https://syngoportal.org/ (bulk download SynGO release “20231201”) and https://syndive.org/#exports (raw data info and enrichment; the union of excitatory and inhibitory cortical synaptic genes was used). The list of genes enriched in perisynaptic astrocytic processes, brain endothelial cells, astrocytic types and glia limitans were obtained from Sakers et al.^82^, Yu et al.^44^, Batiuk et al.^42^ and Hasel et al.^45^, respectively. To identify if a transcript was enriched in neuronal or non-neuronal cell-types, we used the data from Yao et al.^43^ and the function FindAllMarkers in the Seurat package^83^. Only genes that were significantly (p<0.05) enriched in a class were labelled as neuronal or non-neuronal. If a gene was enriched in two classes, the class with the lowest p-value was used. If a gene was enriched in more than two classes, it was labelled as neuronal or non-neuronal only if the enriched subclasses were all neuronal or non-neuronal, otherwise it was labelled as not type-specific. Differentially expressed genes were identified after shrinking the log2FoldChange obtained with the DESeq function using the *apeglm* method^84^, with an adjusted p value of <0.05 and no log2FoldChange threshold. GO analysis was performed with the ‘enrichGÒ function of the clusterProfiler package, selecting all ontologies and using the package org.Mm.eg.db to refer to the mouse genome annotations. The enriched gene list was always compared to the list of all genes detected in that particular dataset. Enrichment results were simplified with the clusterProfiler ‘simplify’ function with a cutoff of 0.5 (Jaccard’s similarity index) and plotted, after selecting the top terms according to number of genes, with the enrichplot function ‘emapplot’ (with layout set to “nicely” and min_edge to 0.15). Edges’ width was adjusted manually. For synaptosome analyses, the unsorted data were only used for the PCA plots. For the comparison with the hippocampal data, the data were obtained from Kaulich et al.^29^ and analyzed with the same thresholds (adjusted p value < 0.05 and no log2FoldChange), always comparing the data of a particular stratum to the whole CA1. For synaptosomes, the tdTom+ fraction of the stratum was directly compared to the tdTom+ fraction of CA1 (no comparison was done within fractions from the same stratum).

## Resource availability

The raw transcriptomic data, code and processed data will be made available on the European Nucleotide Archive (https://www.ebi.ac.uk/ena/browser/home) and on Edmond (https://edmond.mpg.de/) upon acceptance for publication.

## Acknowledgements

We thank Susanne tom Dieck for help with handling animal lines; the MPIBR facilities, in particular the animal facility, imaging facility and mechanical workshop, for their continued support; Beatriz Alvarez-Castelao for advice on metabolic labelling experiments; Marcel Jüngling and Julio Perez for advice on library preparations; Paul Donlin-Asp for assistance with the laser capture sample preparation; Vanessa Stempel and Elena Kutsarova for access to the cryostat; Julia Kuhn for creating graphics of mice, synaptosomes and FASS. The lab of E.M.S. is funded by the Max Planck Society and this work was also supported by the European Union (ERC, DiverseSynapse, 101054512). Views and opinions expressed are, however, those of the author(s) only and do not necessarily reflect those of the EU or the ERC. Neither the EU nor the granting authority can be held responsible for them. E.K. acknowledges funding by EMBO (Postdoctoral Fellowship EMBO ALTF 148-2023).

## Author contributions

T.S. and E.M.S. conceived the project. T.S. and B.N.A. performed the experiments with help from E.K., N.F., L.M., H.W. and E.C.. G.T. preprocessed the transcriptomic data and gave input on the analysis. T.S. analyzed the data and prepared the figures. T.S. and E.M.S. wrote the manuscript.

## Figures and figure legends

**Figure S1.**
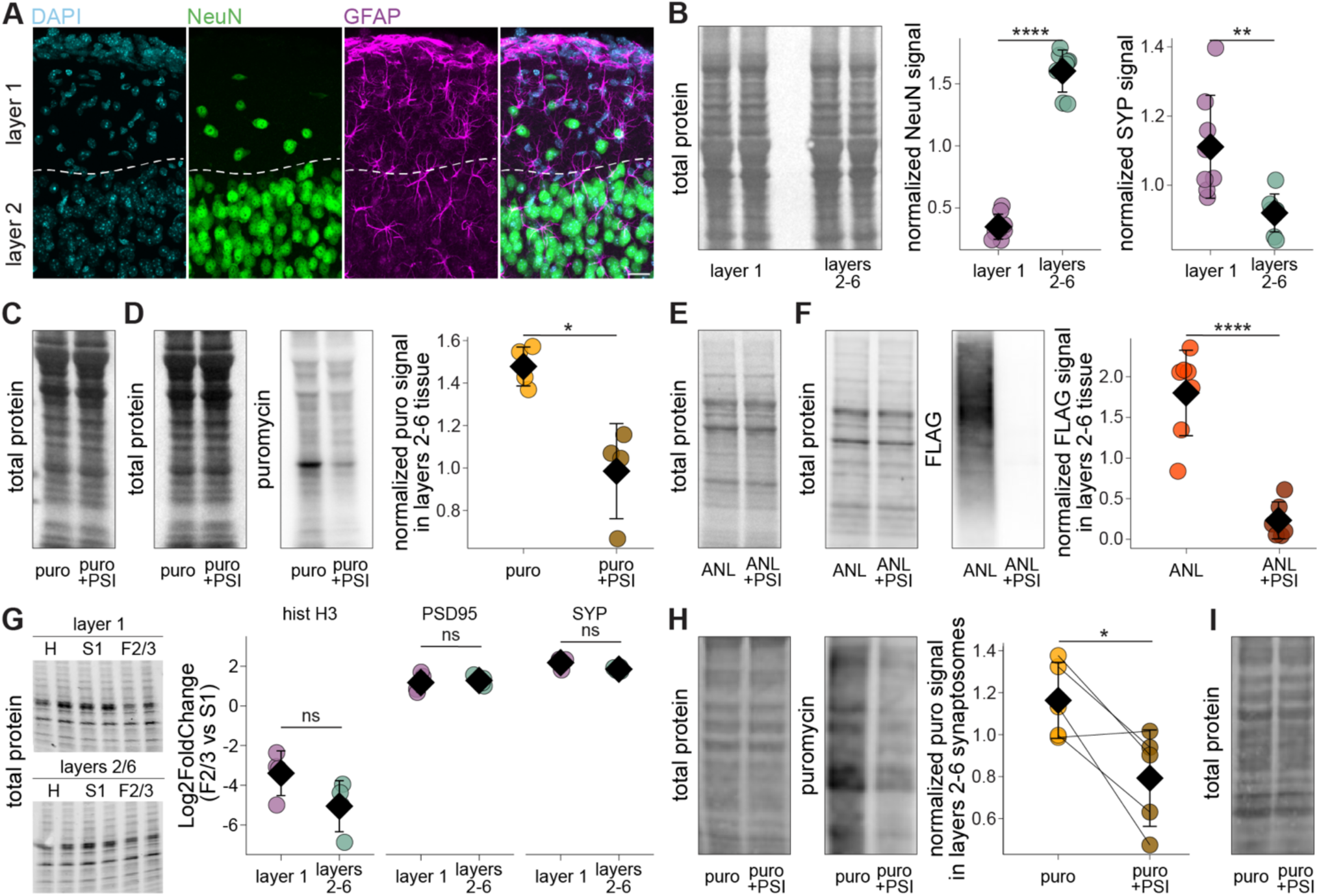
Protein synthesis in neocortical layers 1 and 2-6. Related to Figure 1. A) Superficial layers of the mouse cortex stained with DAPI (cyan), anti-NeuN antibody (green) and anti-GFAP antibody (magenta), revealing a relative paucity of layer 1 cell bodies and those that remain are mostly glial. Scale bar = 20 μm. B) Total protein blot related to main Figure 1E and quantification of NeuN and SYP in layer 1 and layers 2-6 (n = 8 brains, Welch Two Sample t-test). C) Total protein blot related to main Figure 1F. D) Representative Western blot (left) and quantification (right) of protein synthesis in dissected layers 2-6 tissue, labelled in parallel to layer 1 shown in main Figure 1F (n = 4 brains, Welch Two Sample t-test). E) Total protein blot related to main Figure 1G. F) Representative Western blot (left) and quantification (right) of protein synthesis in dissected layers 2-6 tissue from NEX-Cre::3xMetRS* mice, processed in parallel to layer 1 in main Figure 1G (n = 6 or 7 brains, Welch Two Sample t-test). G) Total protein blot related to main Figure 1H and quantification of enrichment (Log2FoldChange) between synaptosome fraction and homogenate after the first centrifugation (S1) of histone H3, PSD95 and SYP in both layer 1 and layers 2-6 (n = 4 brains, Wilcoxon signed-rank test). H) Representative Western blot (left) and quantification (right) of protein synthesis in synaptosomes derived from dissected layers 2-6 tissue, processed in parallel to those from layer 1 in main Figure 1I (synaptosomes split in two conditions, n = 5 brains, paired t-test). I) Total protein blot related to main Figure 1I. * p < 0.05; ** p < 0.01; **** p < 0.0001; ns = not significant. Points represent individual observations; the black diamond and the error bars represent the mean and standard deviation. For statistical details, see Supplementary Table 1.

**Figure S2.**
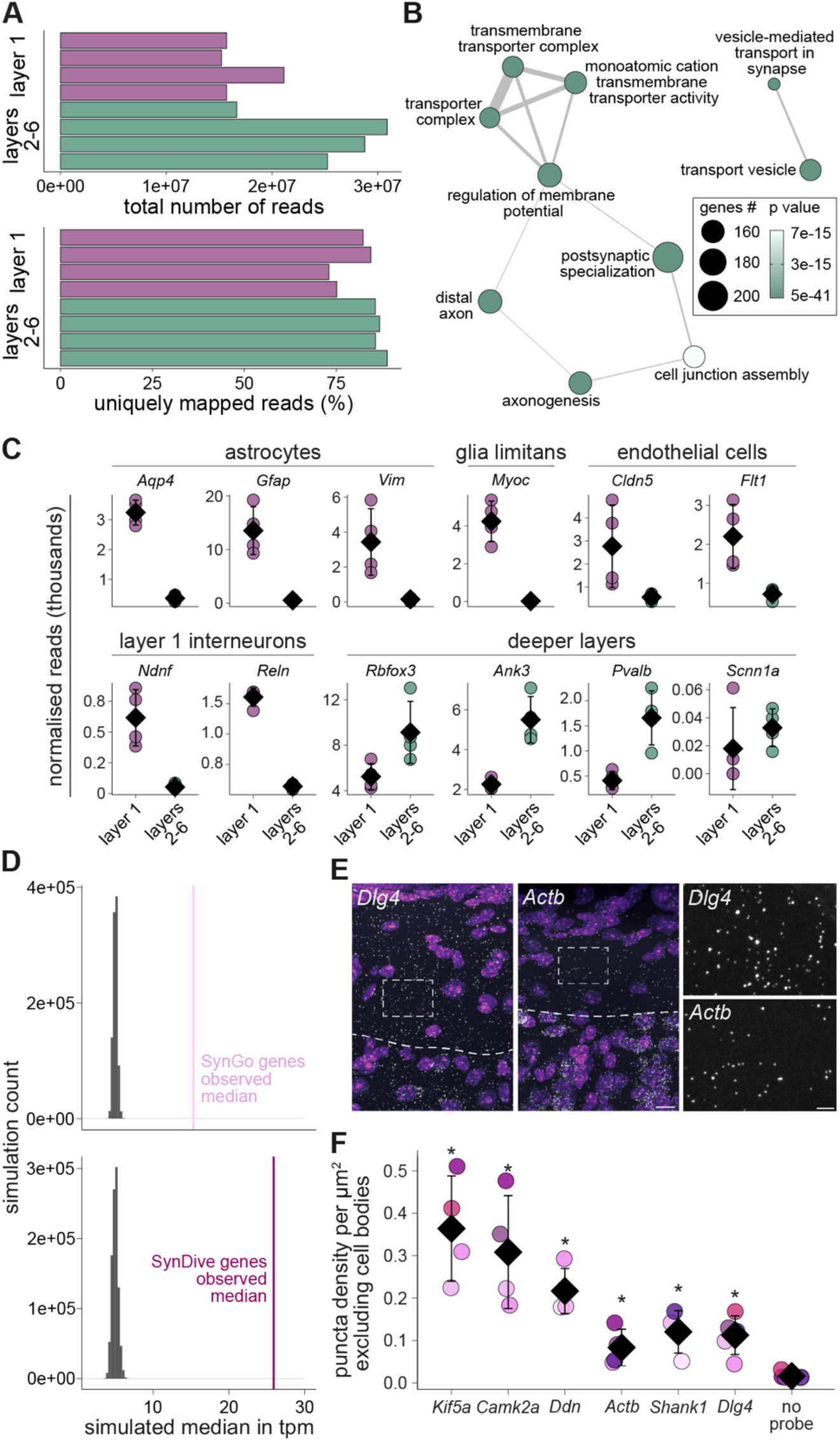
Layer-specific transcriptomes, marker genes and synaptic transcript abundance in layer 1. Related to Figure 2. A) Total number of reads and percentage of reads uniquely mapped to the mouse genome in each library from the laser capture data. B) The 10 most overrepresented GO terms associated with the genes enriched in layers 2-6. The terms are ranked by gene count. The thickness of the edge represents similarity between the terms (only above 0.15). C) Normalized reads in each library for marker genes for layer 1 and deeper layer cell types. DE analysis shows statistically significant differences for all the genes shown. D) Random subsets of genes of the same size as the respective list (SynGO = 1527, SynDive = 783) were drawn 10,000 times, and their median calculated (histograms in gray), which was always lower than the actual median of the lists (thick lines). E) Representative mouse cortex FISH images using *Dlg4* and *Actb* probes (grey), with nuclei stained with DAPI (purple). Dashed lines indicate the layer 1-2 boundary. Dashed rectangles indicate an area devoid of cell bodies, shown at higher magnification on the right. Scale bar on the left and right = 10 μm and 3 μm, respectively. F) Density of FISH puncta in layer 1 after digital exclusion of cell bodies (n = 4 or 5 brains, Welch ANOVA followed by Wilcoxon rank sum exact test – only shown for comparisons against the no probe condition). Points of the same color are derived from data from the same mouse, imaged within the same session. * p < 0.05. Points represent individual observations; the black diamond and the error bars represent the mean and standard deviation. For statistical details, see Supplementary Table 1.

**Figure S3.**
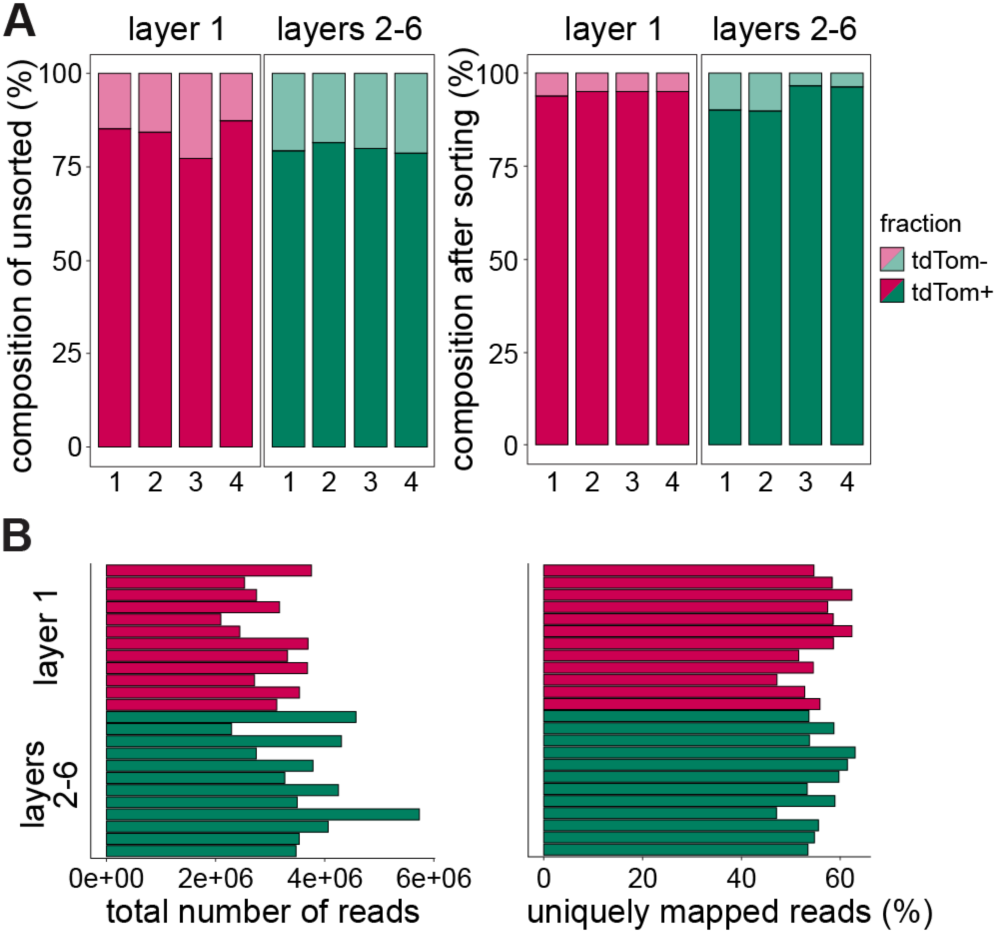
Synaptosome enrichment by sorting and read statistics in NEX-Cre::SypTOM mice. Related to Figure 3. A) Percentages of tdTom+ and tdTom- particles in the unsorted samples (left) and after fluorescence sorting (tdTom+ fraction, right) in layer 1 and layers 2-6 from NEX-Cre::SypTOM mice. B) Total number of reads and percentage of reads uniquely mapped to the mouse genome in each library (one library per fraction, three fractions per layer 1 or layers 2-6 sample).

**Figure S4.**
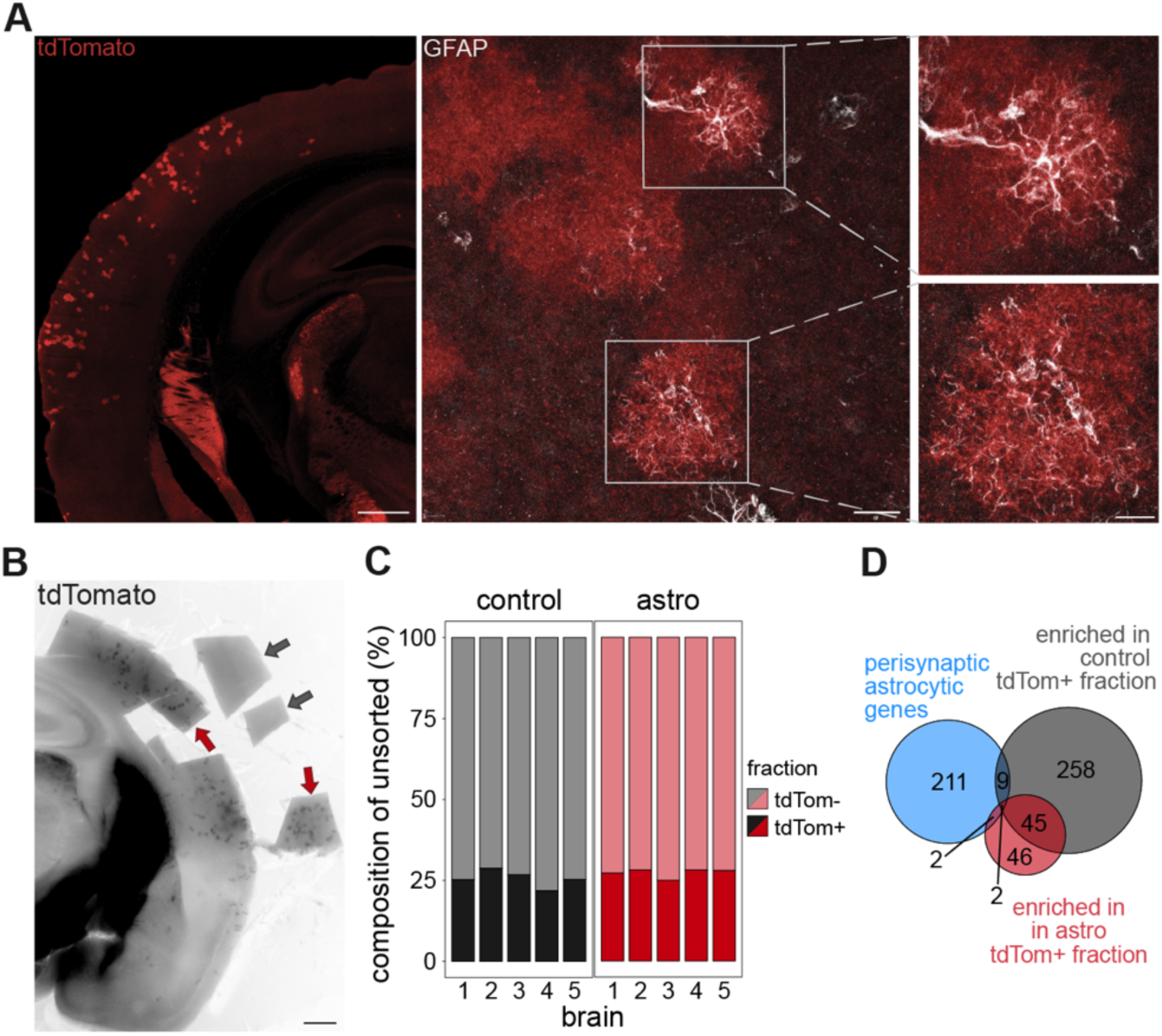
tdTomato-expressing astrocyte minority does not contaminate the sorted synaptosome samples. Related to Figure 4. A) Image of the endogenous tdTomato signal (red) in a coronal slice from a GAD2-Cre::sypTOM mouse (left); higher magnification image on a section stained for GFAP (white, middle) and zoom in on two tdTomato-expressing astrocytes (right). Scale bars from left to right = 500, 20 and 10 μm. B) Representative image of an acute slice after dissection of control samples (gray arrows) and samples enriched in the tdTomato-expressing astrocytes (red arrows). Scale bar = 500 μm. C) Percentages of tdTom+ and tdTom- particles in the unsorted fractions of control (gray) and tdTomato-expressing astrocytes enriched samples (red). D) Venn diagrams showing the overlap of genes found at perisynaptic astrocytic processes^82^ (in blue) with the tdTom+ against tdTom- enriched genes in the control samples (gray) and in the samples enriched in tdTomato-expressing astrocytes (red).

**Figure S5.**
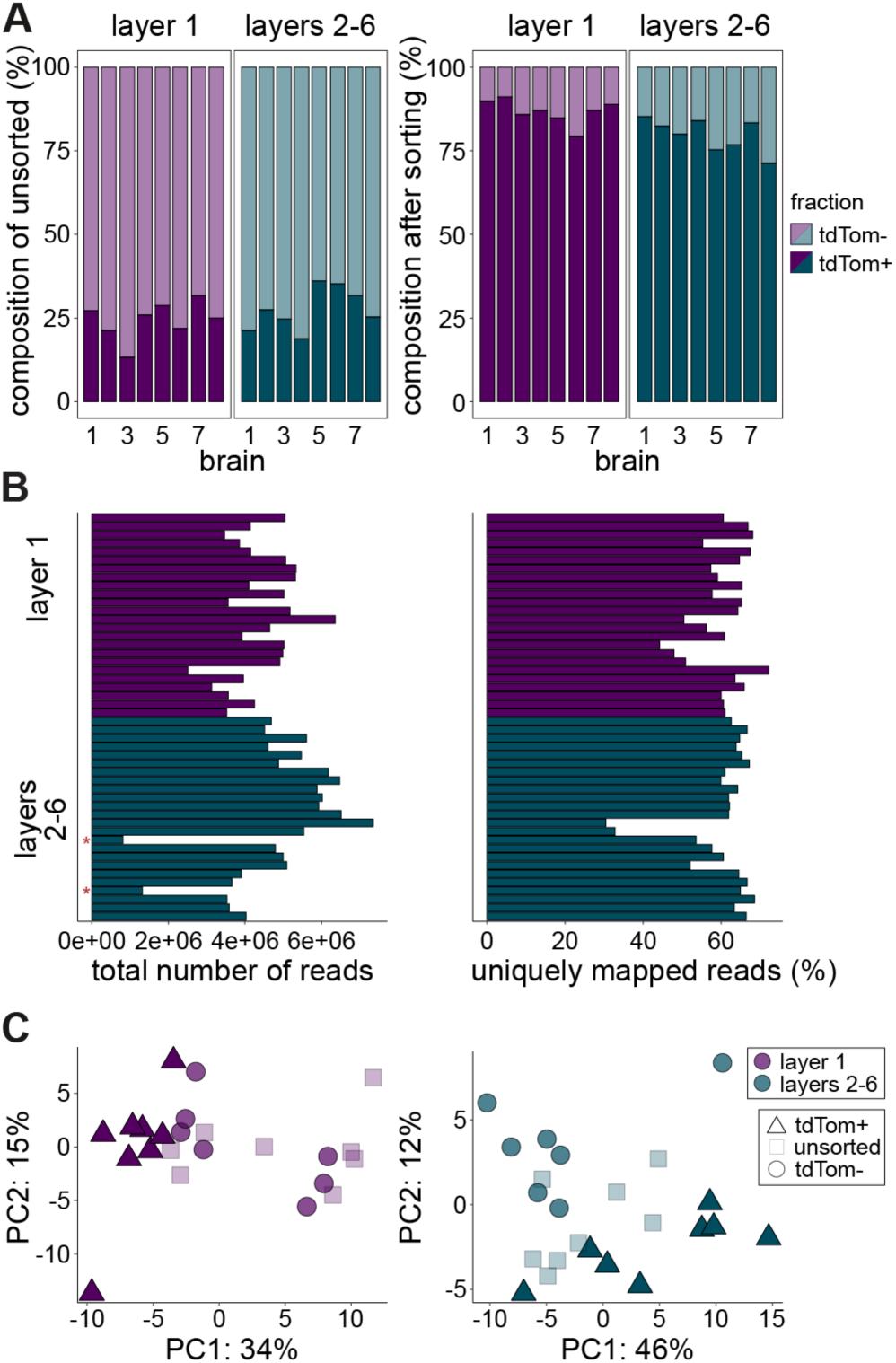
Synaptosome sorting enrichment, read statistics and PCA by layer for inhibitory synapses. Related to Figure 4. A) Percentages of tdTom+ and tdTom- particles in the unsorted samples (left) and after fluorescence sorting (tdTom+ fraction, right) in layer 1 and layers 2-6 from GAD2-Cre::SypTOM mice. B) Total number of reads and percentage of reads uniquely mapped to the mouse genome in each library (one library per fraction, three fractions per layer 1 or layers 2-6 sample). The red asterisks show libraries that were excluded from further analysis. C) First two components in the principal component analysis of the 500 most variable genes after correcting for individual brain variation in the samples from layer 1 and layers 2-6 separately.

**Figure S6.**
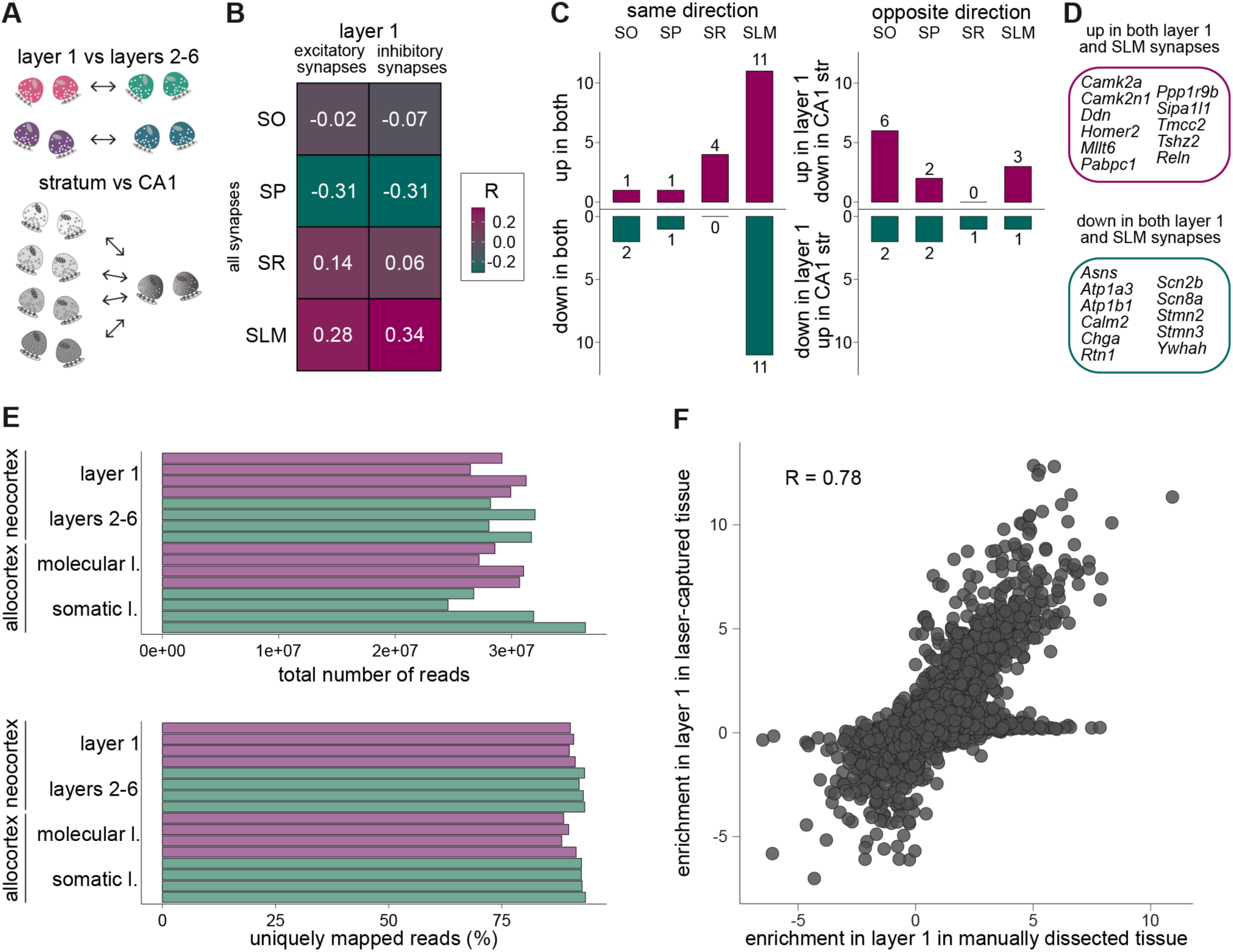
Similarities between layer 1 and SLM at the synaptic level, read statistics for manually dissected tissue and correlation with laser capture data. Related to Figure 6. A) Schematic of the synaptic comparisons used to analyze similarities: excitatory and inhibitory synapses in layer 1 were compared to their layers 2-6 counterparts, all synapses in each CA1 stratum were compared to the whole CA1. Only genes significantly enriched in cortical synapses compared to the negative control particles were kept for the analyses. B) Pearson correlation coefficients between the cortical enrichment in layer 1 excitatory or inhibitory synapses, and the enrichment in synapses from each CA1 stratum. C) Number of genes regulated in the same (left) or opposite direction (right) between layer 1 synapses (union of excitatory and inhibitory enriched genes) and synapses from each CA1 stratum. D) Identity of genes that are enriched or de-enriched at both layer 1 and SLM synapses. E) Total number of reads and percentage of reads uniquely mapped to the mouse genome in each library from manually dissected tissue. F) Correlation of enrichment in neocortical layer 1 in the manually dissected tissue with the enrichment in layer 1 in the laser captured tissue.

---

**Supplementary Table 1.**
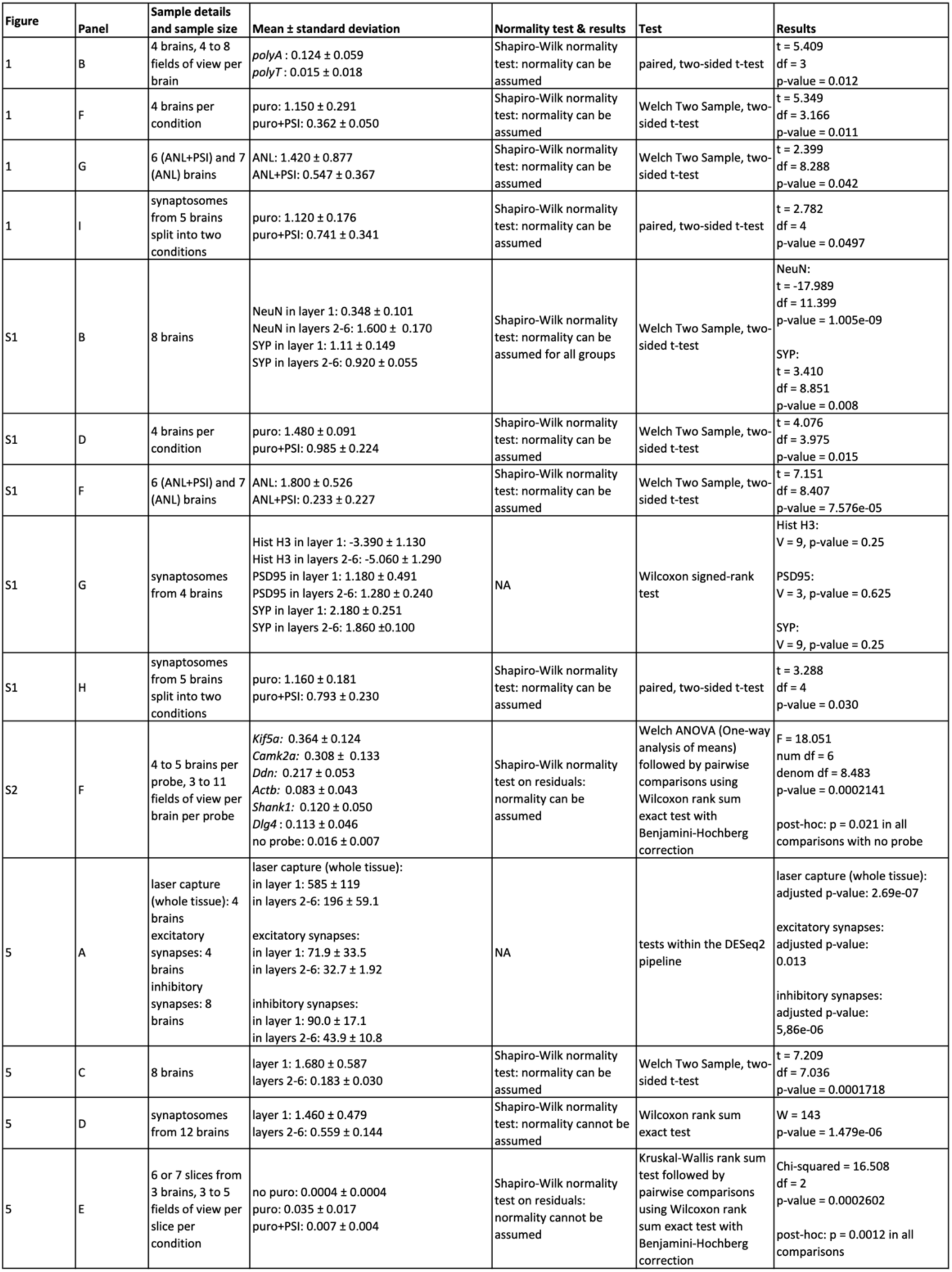
Statistical details.

